# The pangenome of the fungal pathogen *Neonectria neomacrospora*

**DOI:** 10.1101/2021.03.11.434922

**Authors:** Knud Nor Nielsen, Kimmo Sirén, Bent Petersen, Thomas Sicheritz-Pontén, M. Thomas P. Gilbert, Thorfinn Sand Korneliussen, Ole Kim Hansen

## Abstract

The fungal plant pathogen *Neonectria neomacrospora* (C. Booth & Samuels) Mantiri & Samuels (Ascomycota, Hypocreales) is a bark parasite causing twig blight, canker, and in severe cases, dieback in fir (*Abies* spp.). Although often described as a mild pathogen, foresty and phytosanitary agencies have expressed their concern for potential economic impact. Two epidemics caused by this species are known: one from eastern Canada and one current within Northern Europe. We present key genome features of *N. neomacrospora*, to facilitate the research into the biology of this pathogen.

We present the first genome assembly of *N. neomacrospora* as well as the first pangenome within this genus. The reference genome for *N. neomacrospora* is a long-read sequenced Danish isolate, while the pangenome is pieced together using additional 60 short-read sequenced strains covering the known geographical distribution of the species, including Europe, North America, and China.

The gapless reference genome consist of twelve chromosomes sequenced telomere to telomere to a total length of 37.1 Mb. The mitochondrial genome was assembled and circularised with a length of 22 Kb.

The gapless nuclear genome contains a total of 11,291 annotated genes, where 642 only have a hypothetical function, and a 4.3 % repeat content. Two minor chromosomes are enriched in transposable elements, AT content, and effector candidates. Chromosome 12 segregates within the population, indicating an accessory nature. The pangenome compile 15,101 genes, 34% more genes than present in the single isolate reference genome of *N. neomacrospora*. These genes organise into 13,069 homologous clusters, of which 8,316 clusters are present in all analysed strains, 985 are private to single strains.

The British Columbian population branched out before the other populations and are characterized by comparatively larger genomes. The increased genome size can be explained by an expansion of repetitive elements.

The comparative analysis finds a higher number of genes with a signal peptide within *N. neomacrospora* and species within the genus compared to the closely related genera. A species-specific pattern is observed in the carbohydrate-active enzyme repertoire, with a reduced number of polysaccharide lyases, compared to other species within the genus. The CAZymes battery responsible for plant cell wall degradation is similar to that observed in necrotrophic and hemibiotrophic plant pathogenic fungi.

The genome size of *N. neomacrospora* is close to the median size for Ascomycota but is the smallest genome within the *Neonectria* genus. Comparative analysis revealed significant intraspecies genome size differences between populations explained by a difference in repeat content. Isolates with the smallest genomes formed a monophyletic group consisting of all strains from Europe and Quebec. Based on the field observations, we assume that *N. neomacrospora* is a hemibiotroph. Our analysis revealed a secretome consistent with a hemibiotrophic lifestyle.

## INTRODUCTION

The number of complete genome assemblies available for fungi is steadily increasing, and they often have been used as references in species descriptions. Single genomes as a representative of a species can, however, not capture the full genetic diversity within a species and can result in the neglect of crucial structural variation. Describing the full gene repertoire of a species is referred to as the pangenome (McCarthy and Fitzpatrick, 2019), and the first compiled for a eukaryotic pathogen was based on five complete genome assemblies of *Zymoseptoria tritici* in 2018 (Fouché, Plissonneau and Croll, 2018; Plissonneau, Hartmann and Croll, 2018). In essence, the pangenome identifies from the available genomic data across strains the core and accessory genes, based on the presence-absence pattern across the dereplicated genomic features. This framework not only improves the definitions of boundaries or sufficient condition for species delimitation (Doolittle and Zhaxybayeva, 2009), but it can also reveal the biogeographic partitioning within microbial lineages (Reno *et al*., 2009; Porter *et al*., 2017).

Species within the ascomycete genus *Neonectria* are generally found on the bark of recently dead woody plants. Some species can be pathogens causing cankers or twig blight and, in severe cases, dieback of infected trees. Beech bark disease is caused by *N. coccinea, N. faginata* and *N. ditissima*. The latter is also the agent causing European Canker, resulting in substantial economic losses in apple production worldwide, by killing fruit trees and rotting stored fruit (Gómez-Cortecero *et al*., 2016). This pathogen shows a broad host range, which besides *Fagus* and *Malus* also includes genera such as *Acer, Betula, Populus*, and *Salix*. The sister species *Neonectria neomacrospora* (Booth & Samuels) Mantiri & Samuels [anamorph: *Cylindrocarpon cylindroides* var *cylindroides* Wollenw.] has almost exclusively been observed on the softwood genus *Abies* and from the dwarf mistletoe, *Arceuthobium tsugense*. The latter parasitise conifers within the genera *Abies, Pinus* and *Tsuga* (Hawksworth, 1996; Rietman, Shamoun and Van Der Kamp, 2005).

On *Abies, N. neomacrospora* is a bark parasite feeding on the living cells, resulting in heavy resin flow, cankers, twig blight, and dieback. *N. neomacrospora* has been known for more than a century (Wollenweber, 1913) and has historically been perceived as a mild pathogen, mostly attacking already weakened trees (Robak, 1951). *N. neomacrospora* was the likely cause of damages and dieback of *Abies* in Norway in the 1940s (Robak, 1951), but only two epidemic scale outbreaks have been recorded. The first was reported in 1965 on Anticosti Island, eastern Canada, where the majority of the firs were cankered and up to 10% recently dead. Isolates from the Quebec epidemic were at the time compared to isolates from British Columbia (BC) and Norway, and inoculation tests showed that the isolates from the epidemic were more aggressive than isolates from the two other populations, and the existence of races within the species was hypothesised (Ouellette, 1972). The last decade has seen a dramatic increase in damage on fir attributed to *N. neomacrospora* across northern Europe, causing substantial losses to the production of Christmas trees and greenery as well as ornamental trees in urban settings. Different perception of the species exists: the species is considered as a biocontrol agent of mistletoe on conifers in BC (Rietman, Shamoun and Van Der Kamp, 2005), whereas it in Europe is a concern for the phytosanitary authorities (EPPO, 2017).

For woody plants, canker diseases are among the most destructive and difficult to manage problems worldwide (Yin *et al*., 2015). However, plant-pathogen interactions are largely undescribed in forest-ecosystems compared to agricultural systems. This is reflected in the relatively few genomic resources on forest pathogens compared to the resources found on agricultural plant pathogens such as *Magnaporthe, Fusarium, Zymoseptoria, Puccinia*, and *Melampsora* species.

Filamentous fungi can vary tremendously in genome architecture among species, with genome sizes ranging almost three orders of magnitude from a few megabases to gigabase sizes (Stajich, 2017). Third-generation sequencing (TGS), also known as long-read sequencing, has made it possible to sequence across repeat regions that previously fragmented genome assemblies. This was done for the biotrophic barley powdery mildew pathogen *Blumeria graminis*, which had its repeat content corrected from 64% (Spanu *et al*., 2010) to 74% (Frantzeskakis *et al*., 2018), by sequencing trough the large repeat regions using long-read sequencing. Thus, TGS enables complete chromosome assembly, including previously difficult to assemble repeat-rich regions such as centromeres and telomeres (Dallery *et al*., 2017).

Repeat content can accumulate in specific regions or chromosomes (Faino *et al*., 2016) or it can be dispersed throughout the genome as in the case of *B. graminis* (Frantzeskakis *et al*., 2018). The accumulation of repeats and especially transposable elements (TE) have both direct and indirect effects locally on genes. TE can facilitate gene duplications of neighbouring genes, but they are also thought predominantly to disrupt genes by moving into exons. The potentially adverse effects explain why the genome has different ways of inactivating TE’s. A fungal specific, although not a universal method, is the silencing by repeat-induced point mutations (RIP) (Selker *et al*., 1987). RIP will degrade TE by cytosine-to-thymine mutations and guanine-to-adenine mutations, locally reducing the guanine-cytosine (GC) content (Irelan and Selker, 1996). The RIP mechanism is not a precision tool and genes situated near TE are likely to experience an elevated mutation rate as collateral damage, known as RIP leakage. Both repeat activity and RIP are thought to play a role in the evolution of secreted virulence proteins, termed effectors (Dong, Raffaele and Kamoun, 2015; Derbyshire *et al*., 2017). Effectors modulate host defences facilitating infection, and co-evolve with host defences.

Besides effectors, the carbohydrate-active enzymes (CAZyme) play a vital role in the plant pathogenic secretome. They facilitate infections in the appressorial phase and necrotrophy by degrading cell walls (O’Connell *et al*., 2012). Therefore, differences in the enzyme repertoire between fungi can elucidate fungal lifestyle, primary substrate and possible infection routes.

The heterogeneous accumulation of repeat content across the genome observed in some filamentous plant pathogens has become known as the two-speed genome (Croll and McDonald, 2012; Raffaele and Kamoun, 2012). These bipartite genomes have regions undergoing different evolutionary trajectories. A well-described example of this is *Fusarium oxysporum*, which has eleven relatively conserved core chromosomes and then one or more TE rich, highly recombinant and fast-evolving chromosomes which presence or absence determine the pathogens host range (Van Da*m et al*., 2017). Although important in pathogen-host interactions, these repeat-rich chromosomes are expendable and are also termed accessory chromosomes.

The objective of this study is to give an overview of the genetic constitution of *N. neomacrospora*. We report on the gapless genome assembly of this fungal plant pathogen, obtained using Pacific Biosciences SMRT sequencing, resulting in the first available genome of this species. In order to unravel the known geographical distribution of the species, further 60 isolates of *N. neomacrospora* were sequenced. This comprehensive dataset is used for investigating the genetic basis of the putative races, and for compiling a pangenome summarising intraspecies variation. We identify species-specific genes by comparing the *N. neomacrospora* pangenome gene collection to six related species within the genus, and an additional nine species from within the Nectriaceae family.

## MATERIALS AND METHODS

### Strains

The *Neonectria neomacrospora* isolate KNNDK1 was isolated from an *Abies nordmanniana* twig showing the typical symptoms of twig blight: needle loss and bark surface depression. The collection took place on the 22^ed^ of July 2015 at Silkeborg Nordskov (56.1634, 9.5745, ±2m), Denmark.

In addition to strain KNNDK1, 60 strains of *Neonectria neomacrospora* are included in the pangenome analysis. These strains were obtained either by field collections or from revivable herbarium collections. Sixty unique strains, derived from the clone-censuring of a collection of 66 isolates. The collection covered the known geographical distribution of the species, including Europe (n=38), North America (n=21) and China (n=1) (Table S1). Fifteen non-*N. neomacrospora* isolates were included in the study, representing related species within the Nectriaceae family. Five strains, collected in Norway, the Netherlands and France between 1957 and 1961, were obtained from Westerdijk Fungal Biodiversity Institute (CBS), The Netherlands and the Norwegian Institute of Bioeconomy Research, NIBIO. Five strains collected in 1967 from the outbreak centred on Anticosti Island, Quebec was obtained from The René Pomerleau Herbarium, Laurentian Forestry Centre (CFL), Canada. Two isolates from British Columbia from 1996 and 2005 were also obtained from CBS. A single *N. neomacrospora* strain from the Hubei province in China from 2014 was provided by the Herbarium Mycologicum Academiae Sinicae (HMAS). Forty-eight strains were collected between 2015 and 2019. All strains were sampled from individual trees, ensuring that the same individual was not sampled twice. With a few exceptions, all contemporary samples from Europe and North America were geo-referenced when collected (Table S1).

In addition to the cultured samples, six genome assemblies were downloaded from NCBI Genbank (Table S1). These samples added three genera within the family Nectriaceae and the species *Metarhizium anisopliae*, family Clavicipitaceae, as an outgroup to the phylogeny and comparative analysis. Further details on all isolates are given in the supplemental material, Table S1.

### Cultures, DNA extraction and sequencing

Macroconidia were collected from sporodochia on the bark of infected *Abies* sp., using the tip of a needle. When sporodochia were not available, the fungus was isolated from the wood and microconidia were collected from these cultures. Axenic single-spore cultures were derived by plating a small number of conidia diluted in water on potato dextrose agar (PDA) plates, which allowed conidia to separate. After 24 h of incubation, plates were observed under a dissection microscope at 50× magnification and single germinating conidia were collected and transferred to new PDA plates. Single-spore cultures were maintained in 20 % (v/v) glycerol at −80 °C.

### DNA extraction and sequencing

The isolates were transferred to potato dextrose broth (PDB) for 4-5 days at room temperature, and the mycelium was collected on Whatman filter paper (grade 1), rinsed with water and lyophilized. 20-40 mg dried mycelium was homogenized with 200 mg 1 mm zirconia beads in a bead mill (Retsh Mixer Mill MM301) prior to DNA extraction.

Strain KNNDK1 had DNA extracted using a CTAB protocol. Chemical and enzymatic lysis was achieved by 1-hour incubation at 65°C, mixing every 10 min with 550 µl CTAB buffer (2% CTAB, 0,1%M Tris/HCl, 1,4M NaCl, 0,02M EDTA, 1%PVP, 0,1% 2-mercaptoethanol) and 3 µl RNase A (20mg/ml). Subsequently, 550 µl chloroform:isoamyl alcohol (24:1) was added to each sample, gently mixed by inversion and centrifuged for 10 min at ∼18000xg at 4°C, and the aqueous supernatant was transferred to a new 2 ml tube. To the transferred aqueous phase, two volumes (700 µl) of ice-cold 100% ethanol and 200 µl 2M NaCl was added and incubated at -20°C for 2 hours. Samples were then centrifuged for 20 min at ∼18000xg at 4°C, the supernatant removed and the pellet washed twice with 1 ml 70% ethanol and ten mM Ammonium Acetate (a volatile salt to displace the NaCl) by gently inverting the tube and incubation at room temperature for 10 min and then centrifugation 5 min at ∼18000xg. Excess ethanol was removed by pipetting, and the DNA pellet was air-dried for 15 min at 37°C. The pellet was dissolved in a 50% concentration of DC5 MoBio PowerClean kit (EDTA free) elution buffer. The eight samples were pooled and up-concentrated using a speed vac for two hours, condensing 10x to a final conc. of 0.2 µg/µl. The remaining sample had DNA extracted using the DNeasy UltraClean® Microbial DNA Isolation Kit (Qiagen) with the addition of Proteinase K 1% to the lysis mix and a prolonged lysis incubation of 2 hours at 62 °C. DNA was purified with the DNeasy PowerClean Pro Cleanup Kit, and concentrations determined using a Qubit 3 Fluorometer with the Qubit™ dsDNA BR Assay Kit.

From the isolated high-molecular-weight DNA of strain KNNDK1, a 20-kb size-selected library was constructed and sequenced on ten SMRT cells using the PacBio (Pacific Biosystems) RSII instrument with P6-C4 chemistry. Library construction and sequencing were done at the Duke GCB Genome Sequencing Shared Resource (Durham, NC, USA). The remaining DNA extracts were fragmented by sonication to 200-800 bp using the Covaris M220. Sequencing libraries were constructed following the protocol described in Carøe et al. (Carøe et al. 2018), using 100-300 ng dsDNA, and dual-indexed with seven bp Extraction, library and index PCR blanks were included to evaluate the potential contaminations during the library building process. No blanks amplified in qPCR quantification step, and were therefore not sequenced. To ensure library complexity, amplification was done in duplicates and subsequently pooled prior to purification with SPRI-beads. Indexed libraries were quantified on a 5200 Fragment Analyser System (Agilent), and an equimolar pool of all libraries was produced. The pooled library was purification using a BluePippin (Sage Science, Beverly, MA, USA), selecting fragments between 200 bp and 1000 bp. Libraries were sequenced in one lane of an Illumina NovaSeq 6000 SP 150 PE sequencing at the Danish National High-Throughput DNA Sequencing Centre.

### Long-read genome assembly

SMRT reads were assembled using Canu v. 1.3.4 (Koren *et al*., 2017), assuming a genome size of 38 Mb, and further assembled into scaffolds with SSPACE_LongRead v.1.1 (Boetzer and Pirovano, 2014). The genome assembly was improved by gap-filling with PBJelly from the pbsuite v.15.8.24 (English *et al*., 2012). Before and after scaffolding, raw-reads were mapped to the assembly with Minimap2 v.2.6 (Li, 2018), and remaining insertion-deletion and base substitution errors were reduced by polishing the consensus sequence using Arrow v2.3.3 (https://github.com/PacificBiosciences/GenomicConsensus). All assembled nuclear contigs <50 kb in size were discarded.

Identification of the contig corresponding to the mitochondrial genome was facilitated via BLASTn searches using the assembled contigs as queries against the assembled mitochondrial genome of *Fusarium oxysporum* strain F11 (GenBank AY945289.1; Pantou, Kouvelis and Typas, 2008). The contig was then circularized and trimmed using the program Circlator (Hunt *et al*., 2015). Telomeres were identified at chromosomal ends as the (5′-TTAGGG-3′)n simple sequence repeats characteristic for filamentous fungi (Wu *et al*., 2009). This was done using RepeatMasker v4.0.7 (see URLs).

### Short-read genome assembly

Illumina reads were trimmed, removing adapter and primer sequences and bases at read ends with Phred quality below 20 (-q20), while only keeping reads longer than 80 bp. This was performed using AdapterRemoval (v.2.2.4)(Schubert, Lindgreen and Orlando, 2016), options: [--trimns --trimqualities --minquality 20 --minlength 80]. Trimmed reads were de-novo assembled using SPAdes v.3.13.1 (Bankevich et al. 2012) (kmers 21, 33, 55, 77, 99, 127) using mismatch and short indel correction with the Burrow-Wheeler Aligner, BWA-MEM v.0.7.12 (Li 2013). Assemblies were improved using Pilon v.1.22 (Walker et al. 2014). The assembly summary statistics were calculated using Quast v5.0.2 (Mikheenko *et al*., 2018) (Table S2).

### Annotation

A custom repeat library was created by adding *de novo* identified repeats from RepeatModeler (Hubley et al., 2016) to the repeat databases, Dfam_Consensus-20170127 (Hubley et al., 2016) and RepBase-20170127 (W. Bao et al., 2015). RepeatModeler was run on the genome masked by these two public databases, and the iteratively growing custom library, using RepeatMasker v4.0.7 (Smit et al., 2015) with the option [-species] set to fungi. RepeatModeler relies on the three *de novo* repeat finding programs RECON (Z. Bao & Eddy, 2002), RepeatScout (Price et al., 2005) and LtrHarvester/Ltr_retriever (Ellinghaus et al., 2008). Repeats were identified using RepeatMasker [option: sensitive] and the generated custom repeat library. The custom library was built on the following six *N. neomacrospora* isolates: two European (KNNDK1, CH01011), two from British Columbia (CA03011, CBS 118984), a single Quebec isolate (CA01061) and the Chinese isolate (HMAS 252906) (Isolate metadata: table S2).

Gene prediction and functional annotation of the polished assembly was conducted using the Funannotate pipeline v. 1.6.0, (https://github.com/nextgenusfs/funannotate). Repeats were identified with RepeatModeler and soft masked using RepeatMasker. Protein evidence from a UniProtKB/Swiss-Prot-curated database (downloaded on 2019-08-08) (Bateman *et al*., 2017) was aligned to the genomes using tBLASTn and Exonerate (Slater and Birney, 2005). Two gene prediction tools were used: AUGUSTUS v3.2.3 (Stanke and Morgenstern, 2005) and GeneMark-ES v4.32 (Besemer and Borodovsky, 2005), with *Fusarium graminearum* as the model for the AUGUSTUS gene predicter and BAKER1 (Hoff *et al*., 2016) for the training of GeneMark-ES. tRNAs were predicted with tRNAscan-SE (Lowe and Eddy, 1997). Consensus gene models were found with EvidenceModeler (Haas *et al*., 2008). Functional annotations for the predicted proteins were obtained using BLASTP to search the UniProt/SwissProt protein database (2019-12-11). Protein families (Pfam) and Gene Ontology (GO) terms were assigned with InterProScan 5.36-75.0 (Jones *et al*., 2014). The secretome was predicted using SignalP v5.0 (Almagro Armenteros *et al*., 2019), identifying proteins carrying a signal peptide. To further characterize the secretome, putative CAZymes were identified using HMMER v.3.2.1 (Eddy, 2011) and family-specific hidden Markov model (HMM) profiles of dbCAN2 meta server (Lombard *et al*., 2014). Putative proteases and protease inhibitors were predicted using the MEROPS database (2017-10-04) (Rawlings, Barrett and Finn, 2016). All functional predictions and annotations were executed through the Funannotate pipeline (see URLs). The completeness of the assembled genomes and the corresponding gene annotations was evaluated using the single-copy ortholog genes from Ascomycota dataset of BUSCO2 (Simão *et al*., 2015).

### Genome-wide analyses

GC content was calculated in 10 kb windows using seqkit v.0.7.1 (Shen *et al*., 2016). Sequence coverage was calculated using the SAMtools v.1.9 depth command (Li *et al*., 2009). Gene density was calculated in 10 kb windows using the R package RIdeogram v.0.2.2 (Hao *et al*., 2019), and visualised together with GC content using the same package.

### Clustering

The predicted proteomes were used for pangenomic analysis following the protocol proposed and discussed by Eren et al. (2015) and Delmont and Eren (2018). Briefly, the amino acid sequences of the genes underwent an all against all search using BLASTP 2.2.31 (Altschul *et al*., 1990). Using the resulting bit-score, we calculated the minbit heuristic (Benedict *et al*., 2014), and retained all the pairwise edges with a higher than 0.5 minbit-score. The resulting edge list with the per cent identity was fed into MCL v14.137 (Van Dongen and Abreu-Goodger, 2012) for clustering with default parameters, except inflation, where we increased to 10. Multiple sequence alignments (MSA) were generated from the gene cluster members using Muscle v3.8.1551 (Edgar, 2004). If a gene cluster had genes from all the samples, we denoted those clusters as core gene clusters. Furthermore, if the genes were found as a single copy, we defined those clusters as single-copy core gene clusters.

Similarly, species-specific gene groups were identified by the same presence-absence strategy. In total, this strategy was used to resolve the family, genus, and species level unique gene clusters as well as the sub-population gene clusters for *N. neomacrospora*. Rarefaction curves were generated for the count of core (shared by all), accessory (shared by some), and singleton gene cluster. Strains were randomly drawn without replacement, and pangenome statistics are based on one hundred random iterations.

EggNOG-mapper v2.0.1 (Huerta-Cepas *et al*., 2017) was used to annotate all the genes to functional categories using the eggNOG database v.5.0 (Huerta-Cepas *et al*., 2019) and DIAMOND v0.9.24 (Buchfink, Xie and Huson, 2015). SignalP version 5.0b (Almagro Armenteros *et al*., 2019) was used to annotate the signal proteins. The dbCAN annotation tool for automated CAZyme annotation, Run-dbcan v2.0.3 (H. Zhang *et al*., 2018), was used with default parameters to predict CAZymes using CAZyDB version 07312018. We further added: 1. ‘Predicted taxonomic group’, ‘Predicted protein name’, ‘COG Functional Category’ and ‘eggNOG free text description’ from eggNOG-mapper output, 2. Prediction and the ‘CS Position’ from SignalP, and 3. HMMER v3.1b2 (Eddy, 2011), Hotpep (Busk *et al*., 2017), DIAMOND v0.9.24 (Buchfink, Xie and Huson, 2015) predictions from CAZymes annotations of single genes to the gene clusters, by assigning the results to the gene cluster if more than 90 % of the member genes had the same function. If lower, we assigned the highest match as hypothetical with the proportion indicated in the name. Throughout the analysis, all the additional data parsing and processing were done using Jupyter notebooks (Kluyver *et al*., 2016) and python3 unless stated otherwise. Hierarchical clustering using the R Stats package function ‘hclust’, with 8euclidean distances and complete linkage was made of all strains within the study by their complete CAZyme profiles. CAZymes were grouped based on the substrates as listed by Rytioja *et al*., (2014) and clustered as described above based on their abundance across all studied strains. Clustering and visualization of CAZyme profiles were done using the R package pheatmap v.1.0.12 (Kolde, 2018).

### Anploidy

The breadth of sequence coverage of the 60 isolates of *N. neomacrospora* relative to the reference genome was investigated by summarising bam files using Samtools v.1.9 (Li *et al*., 2009). The breadth of coverage is here defined as how many bases of the long-read reference are covered by the short-read samples at a level of 1X or higher. Coverages of chr11 and chr12 were compared to the mean coverage of the remaining chromosomes (chr1 to chr10), Figure S2.

### Phylogeny

Clustering revealed 1,418 single-copy gene clusters. Transcriptomes of these gene clusters were aligned using MAFFT v.7.402 [option: linsi] (Katoh and Standley, 2013), and filtered on amount of gaps and inter-gene distance, leaving 51 genes with less than two per cent gaps and a minimum inter-gene range of 10kb (on the reference genome KNNDK1). Table S3 lists the annotations of the 51 genes on strain KNNDK1. Substitution models for each codon position in each gene were predicted using ModelFinder (Kalyaanamoorthy *et al*., 2017) as implemented in IQtree v.1.6.12, and used with the concatenated protein alignment to generate a consensus maximum likelihood phylogeny based on 100 trees. The consensus tree was subsequently validated with 100 bootstrap replicates using IQtree v.1.6.12 (Nguyen *et al*., 2015; Chernomor, von Haeseler and Minh, 2016). The outgroup *Metarhizium anisopliae*, family Clavicipitaceae, is not shown in the phylogeny.

### Statistical analyses

The degree of correlation between the presence of transposable elements and GC-content was analysed by linear regression. Principal component analysis on isolate repeat content was make on normalized values of repeat content per repeat caterogy as defined by RepeatMasker. The analysis was performed and visualized in R using the packages ggpubr v.0.2.5 (see URLs) and ggplot2 v.3.2.1 (Wickham, 2016). The difference in mean genome size between populations was tested using a Welch Two Sample t-test. Differences between a single observation and a mean was tested for significance using One-Sample t-test. This is done in exploring differences between of chr11 and chr12 to the mean of chr1-10, summarized in table 1 and table S4.

**Table 1.** Reference genome.

| Chr | Whole-genome | Chr 1-10<br>mean | sd | Chr 11 |  | Chr 12 |  |
| --- | --- | --- | --- | --- | --- | --- | --- |
| Size (Mb) | 37.09 | 3.42 | ± 1.58 | 0.87 |  | 0.30 |  |
| GC-content, % | 53.49 | 53.27 | ± 0.95 | 48.47 | *** | 39.12 | *** |
| Number of protein-coding genes | 11,291 |  |  |  |  |  |  |
| Total gene length (Mba) | 18.87 |  |  |  |  |  |  |
| Gene content, % | 52.53 | 52.63 | ± 6.93 | 48.97 |  | 15.91 | *** |
| Gene count, Mb<sup>-1</sup> | 304 | 308 | ± 13 | 308 |  | 123 | *** |
| Species-specific genes, Mb<sup>-1</sup> |  | 16 | ± 4 | 40 | *** | 78 | *** |
| Gene length, bp | 1,645 | 1,656 | ± 87 | 1581 |  | 1274 | *** |
| Proportion of genes with unknown function, % |  | 5.38 | ± 1.05 | 8.21 | *** | 54.05 | *** |
| Number of exons | 33,144 |  |  |  |  |  |  |
| Number of introns | 21,688 |  |  |  |  |  |  |
| Mean number of intron per gene | 1.89 |  |  |  |  |  |  |
| Number of tRNA | 165 |  |  |  |  |  |  |
| Proteases<sup>1</sup>, Mb<sup>-1</sup> | 427 | 11.17 | ± 1.79 | 13.87 | ** | 3.37 | *** |
| CAZymes<sup>2</sup>, Mb<sup>-1</sup> | 504 | 13.45 | ± 4.09 | 20.80 | *** | 0 | *** |
| Genes with signal peptide<sup>3</sup>, Mb<sup>-1</sup> | 26.60 | 27.80 | ± 7.47 | 49.43 | ** | 0 | *** |
| **Repeats<sup>4</sup>, given as % of sequence length** |  |  |  |  |  |  |  |
| Transposable elements | 1.47 | 1.51 | ± 0.88 | 5.86 | *** | 10.49 | *** |
| Retroelements | 1.0 |  |  |  |  |  |  |
| LINE | 0.03 | 0.02 | ± 0.02 | 0 |  | 0 |  |
| LTR | 0.97 | 1.03 | ± 0.81 | 4.37 | *** | 7.47 | *** |
| DNA transposons | 0.07 | 0.07 | ± 0.08 | 0.11 |  | 0 |  |
| Unclassified | 0.40 | 0.39 | ± 0.16 | 1.37 | *** | 3.02 | *** |
| Small RNA | 0.03 |  |  |  |  |  |  |
| Satellites | 0.02 |  |  |  |  |  |  |
| Simple repeats | 2.48 | 2.52 | ± 0.84 | 2.96 |  | 12.73 | *** |
| Low complexity | 0.58 | 0.63 | ± 0.36 | 0.40 |  | 4.19 | *** |
| Total repeat content | 4.58 | 4.70 | ± 2.00 | 9.22 | *** | 27.44 | *** |

Asterisks indicate that data for the mini-chromosomes differ significantly from the mean for chromosomes 1-10 (One-Sample t-test, \*\*\* P <0.001; \*\* P <0.01; \* P <0.05).
<sup>1</sup> Annotated with MEROPS,
<sup>2</sup> Annotated using dbCAN,
<sup>3</sup> Signal peptides predicted with SignalP-5. This group overlap with Proteases and CAZymes (Table S4)
<sup>4</sup> Annotated with Dfam and RepBase databases using RepeatMasker

## RESULTS

### The complete genome assembly of *N. neomacrospora*

High molecular weight genomic DNA was isolated from an axenic monospore culture of *Neonectria neomacrospora* and sequenced on ten PacBio RSII SMRT cells which produced 9.8 Gb of raw sequence reads. After initial quality assessment and filtering, adapter-removal, error correction and trimming, the input sequencing data for genome assembly was reduced four-fold to 250 thousand reads with a mean length of 9.9 kb, approximately 65x coverage. Assembly resulted in 25 contigs. Five contigs were identified as mitochondrial based on the blast results against the mitochondrial genome of *Fusarium oxysporum* strain F11 (GenBank accession AY945289, Pantou, Kouvelis and Typas, 2008). Only one mitochondrial contig could be circularized using Circlator, the remaining four contigs could be mapped to the circularized genome and were discarded as redundant. Four contigs, equaling 0.14% of the final genome size, could be mapped to subtelomeric regions of larger contigs and were discarded. Fourteen contigs were merged into 12 scaffolds with SSPACE-LongRead, Gaps were filled using PBJelly, and the final assembly was polished to remove erroneous base calls and insertions/deletions (indels) using Arrow as described in materials and methods. Two contigs, of 27 kb and 21 kb (0.13% of the genome size), were not merged with larger contigs by the scaffold builder SSPACE-LongRead and were subsequently discarded.

Twelve gapless chromosomes spanning telomere to telomere were assembled, with a total genome size of 37.1 Mb (Table 1). Telomeres consisted of short sequence repeats of (5′-TTAGGG-3′)n spanning between 107 bp and 142 bp with a mean of 126 bp. The mitochondrial genome was assembled into a single contig and circularized with a full length of 22 Kb.

The overall GC content of the reference genome was 53.49%, with the mean GC content of chromosome 1-10 at 53.27% (sd: 0.95) (Table 1). The two minor chromosomes 11 and 12 have lower GC contents, 48.47%, and 39.12%, respectively. GC content was calculated on the entire chromosomes, without masking centromeric and telomeric regions. Given that these genomic elements are known to be characterized by low GC content and it is expected to have a relatively uniform size across chromosomes, they will have a proportionally significant effect on the calculated GC content of minor chromosomes.

GeneMark-ES found 11,203 gene models where Augustus found 10,824 gene models. EvidenceModeler combined the two sets of models and returned a combined set of 11,378 gene models. Eighty-six transposable elements and one short model were excluded, leaving 11,291 gene models (Table 1), where 642 only have a hypothetical function. With a mean gene length across the genome of 1,645 bp, this means that 52.5% of the genome is comprised of a mix of exons and introns, with a mean number of 1.89 introns per gene. Except for chr12, the gene density across the genome with 308 (sd: 13) genes per Mb. Chromosome 12 diverges from the other chromosomes by a significantly lower gene density of 123 genes per Mb and significantly elevated density of simple repeats (both, One-sample t-test, P < 0.01) (Table 1).

The genome-wide repeat content was 4.3% and is dominated by simple repeats, transposable elements and low complexity sequence in the named order. Twenty-seven per cent of the TE are long terminal repeat (LTR) retroelements. However, with an average length of 636 bp, these are longer than the annotated DNA transposons and unclassified TE and constitute 68% of the TE sequence length. The opposite goes for the unclassified TE, which stands for 67% of the annotated TE, but with an average length of just 205 bp, this only summarizes to 27% of the total repeat sequence length (Table 1).

The two minor chromosomes, chr11 and chr12, diverge from the larger chromosomes in terms of significantly higher repeat content, 9.2% and 27.4%, respectively, mainly consisting of simple repeats and retrotransposons (Table 1). TE density is negatively related with GC content (R^2^ = 0.17, P = 1.4×10^−146^, Figure S1), and the two chromosomes have also significantly lower GC content than the average for the ten larger chromosomes (One-Sample t-test, P <0.001). Gene density, TE density and GC content across the genome are illustrated in figure 1. The gene density of chromosome 11 is similar to that of the larger chromosomes but rich in secreted proteins, with one-third of the chromosome consisting of genes coding for proteases and CAZymes, and close to half being genes coding for proteins with a signal peptide (some of which are proteases and CAZymes). Chr12 has only 15.9% of the sequence length occupied by genes, significantly less than the remaining genome; 25 of the 37 genes had no homolog in the consulted annotation databases and are classified as unknown. Both chr11 and chr12 have significantly more unknown genes than the larger chromosomes (One-Sample t-test, P <0.001). The unknown genes of chr12 did not harbour signal peptides and had a size range from 220 to 2,144 bp. Out of 37 genes, twenty were species-specific.

**Figure 1.**
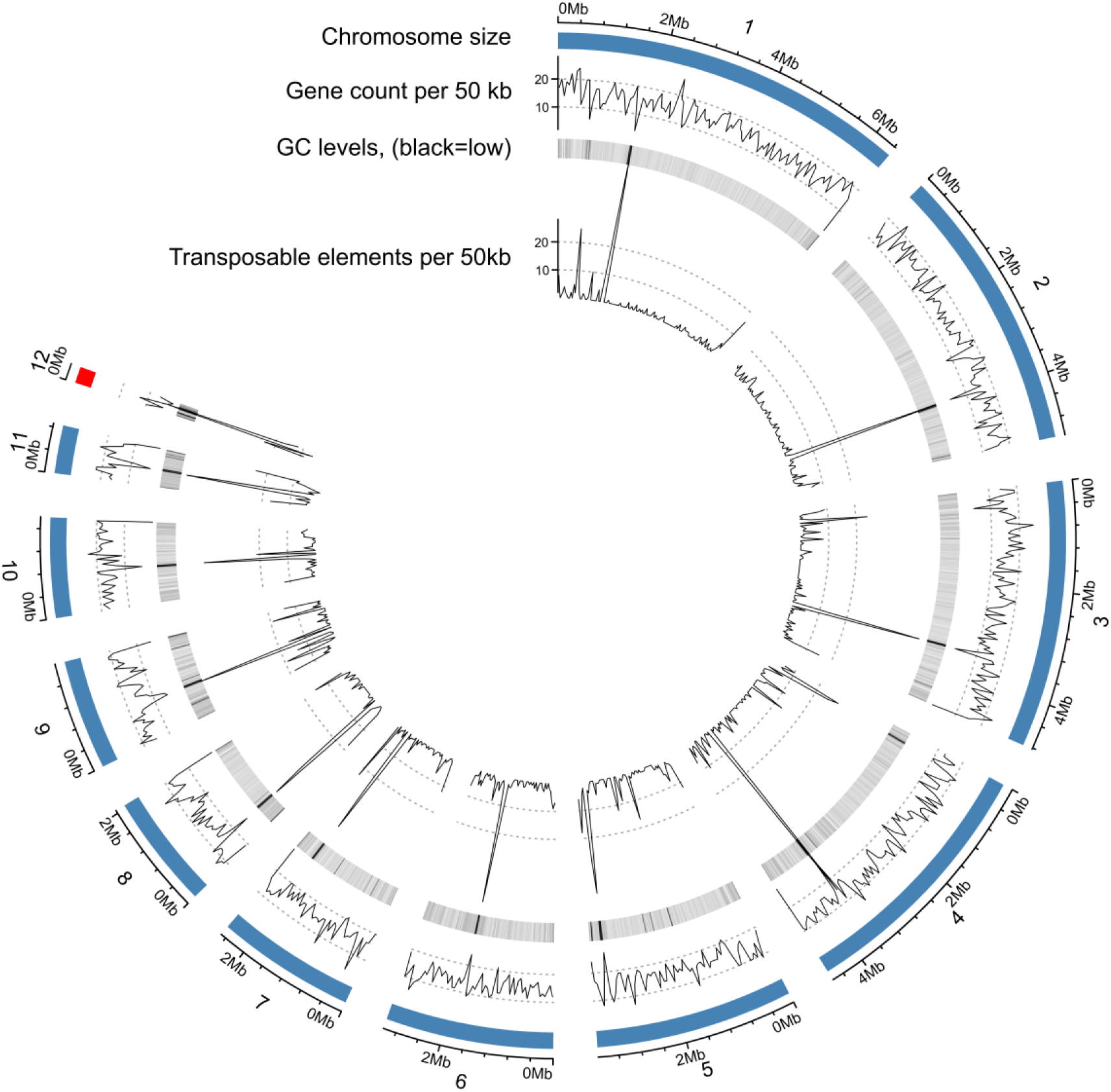
Genome summary of Neonectria neomacrospora strain KNNDK1. Genome-wide statistics of i) chromosome cont and size, ii) gene density, iii) GC level (heatmap with low GC levels in black), and iv) density of transposable elements. Transposable elements and gene is counted in 50 kb windows, where GC is summarized in 10 kb windows. Centromeres are comprised by repetitive elements and have low GC content; each chromosome has a clear centromere candidate region. Chromosome sizes range from 6.4 Mb to 0.3 Mb (for chromosome details, see Table S3).

### Pangenome

We constructed a global pangenome of *N. neomacrospora* based on 61 strains isolated from three continents, spanning the known geographical distribution of the species. The isolates were collected between 1957 and 2019. The 61 strains include the strain KNNDK1 sequenced with TGS as described above and 60 isolates sequenced with Illumina NovaSeq, 150 bp pair-end libraries to a median and average depth of 27x and 30x coverage, respectively (Table S2).

The twelve chromosomes of the reference KNNDK1 were recovered in all but three isolates. The smallest chromosome, chr12 were missing in two isolates from the same location on Anticosti Island collected in 1967 (QFB 19253, QFB 19262) and from one contemporary Norwegian isolate (NO 252140). The proportion of the reference genome covered by reads of the QC and EU strains was lowest at chr11 and chr12. The coverage breadth to the reference for chr1-10 across EU and QC sample range from 90-97% with a mean of 95%. Chr11 ranged from 86-95% with a mean of 91%. Mapping the reads of isolate QFB 19253, QFB 19262 and NO 252140 to the reference genome chr12 only returned a coverage of 4-6% (Figure 2). Chr12 had a relative poor coverage across all isolates, with a mean coverage breadth of 65% (range: 56-70%) and 86% (range: 60-97%) in QC and EU, respectively. The mean coverage breadth of the BC and CN isolates to the Danish reference ranged from 90-92% for chr1-10, 88-90% for chr11, reflecting the greater phylogenetic distance of these sample to the reference. The coverage of BC chr12 ranged from 36-40%, where the CN chr12 had a coverage of 80% more similar to the European samples. The coverage breadth to the reference of chr11 was in all cases slightly lower than the coverage to the ten larger chromosomes. Given the discontinuity in the range of the coverage of chr12, with no other samples from QC and EU having a coverage below 56%, we conclude that the three isolates showing low read coverage to the reference chr12, had lost their chr12, and only have chr1-11. The 4-6% coverage observed could be explained by sequence similarity to other parts of the genome, but this has not been investigated.

**Figure 2.**
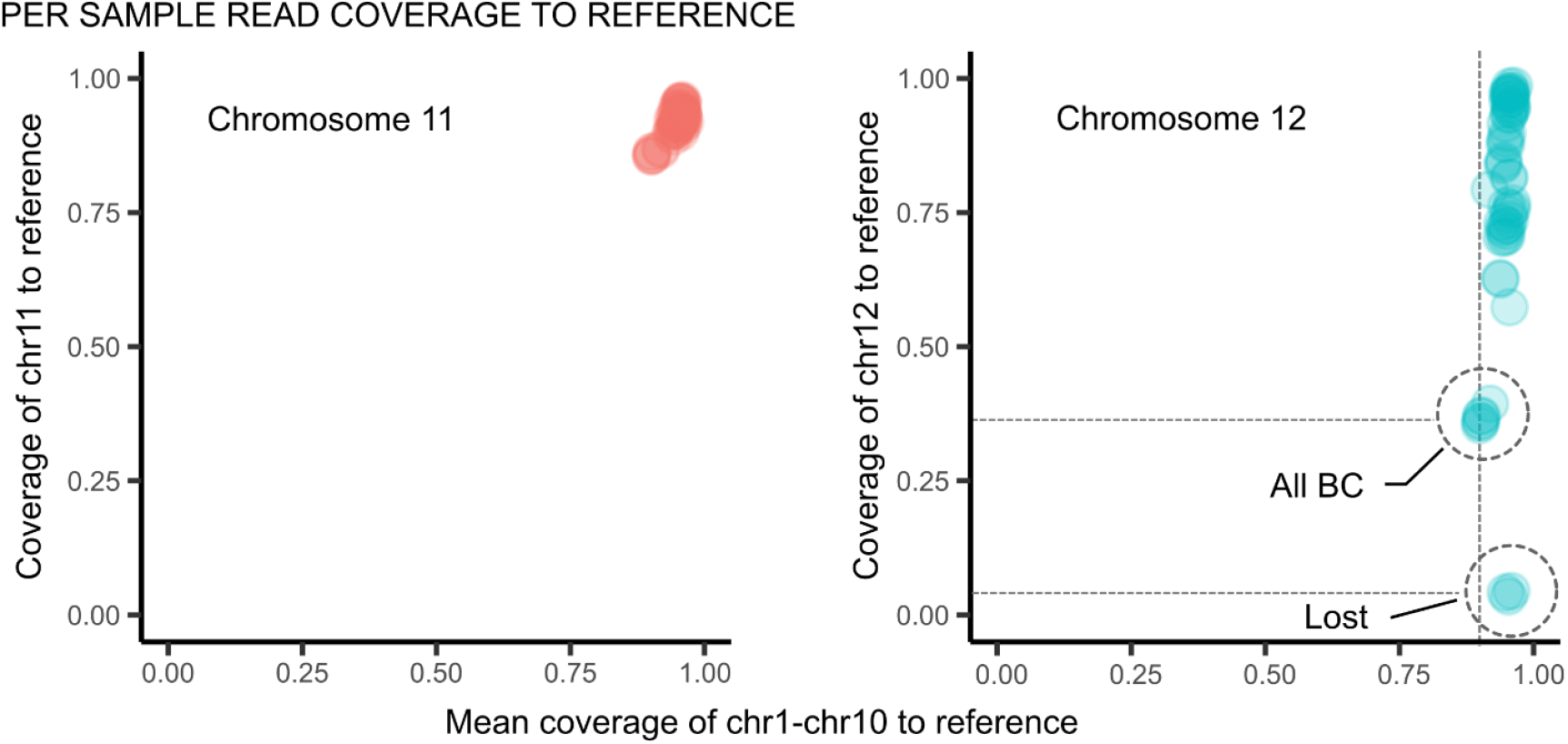
The coverage of chr11 and chr12 relative to chr1-10, each given as the coverage breadth relative to the reference strain KNNDK1. A value of 1 indicate that the reads of the isolate covers the full length of the refence choromosome. Sequencing reads of 60 isolates of *Neonectria neomacrospora* was mapped to the genome of the reference strain KNNDK1. Each dot represent a strain. The x-axis is the mean coverage across the chr1 to chr10 and the y-axis is the coverage of chr11 and chr12. All BC: all six isolates analysed from British Columbia, Canada. Lost: three isolates, two Norwegian and one from Quebec, Canada, with a mean coverage of chr1-10 above 0.9, but a coverage of chr12 below 0.05.

Based on the apparent presence-absence segregation of chr12, its high repeat content and a large proportion of unknown genes (Table 1) we categorize this chromosome as an accessory chromosome.

Significant genome size differences are observed among isolates (Table 2). Genome sizes coincide with the geographical origin of the isolates. BC and China have significantly larger genomes compared to isolates from EU and Quebec, with a mean size of 40.2Mb (sd: 1.3) and 36.9 Mb (sd: 0.4), respectively (Welch Two Sample t-test: P <0.001).

**Table 2.** Features of *Neonectria neomacrospora* genome assemblies. Presented by the single strain KNNDK1 reference, the combined Quebec and European population (QC-EU), the combined British Columbian and Chinese population (BC-CN) and finally by the derived pangenome

| Assembly | Reference | QC-EU sample |  | BC-CN sample |  | Pangenome |
| --- | --- | --- | --- | --- | --- | --- |
|  |  | mean | sd<sup>‡</sup> | mean | sd |  |
| Genome assembly size, Mb | 37.09 | 36.90 | 0.42 *** | 40.18 | 1.29 |  |
| Number of Chr / contigs | 12 | 1,574 | 1,662 | 6,483 | 2,230 |  |
| N50, kb | 4,617 | 176 | 74 | 71 | 18 |  |
| L50 | 4 | 137 | 320 | 172 | 35 |  |
| GC content | 53.49 | 53.81 | 0.31 | 52.14 | 0.98 |  |
| Number of genes | 11,291 | 11,201 |  | 11,234 |  | 15,101<sup>a</sup> |
| Gene clusters | 10,503 | 11,820<sup>b</sup> |  | 11,863<sup>b</sup> |  | 13,069 |
| Core gene clusters |  | 8,073<sup>b</sup> |  | 9,311<sup>b</sup> |  | 8,316 |
| Core genes | 8,436 |  |  |  |  |  |
| Signal peptides | 987 | 975 | 16 | 989 | 21 | 1,419 |
| CAZymes | 427 | 425.8 | 2.3 | 425.6 | 1.5 | 457 |
| Proteases | 395 |  |  |  |  |  |
| CAZymes-Proteases overlap | 28 |  |  |  |  |  |

<sup>‡</sup> sd: one standard deviation
Asterisks indicate that the mean of the two groups differ significantly (Welch Two Sample t-test, \*\*\* P <0.001; \*\* P <0.01; \* P <0.05)
<sup>a</sup> Sum of the highest gene-copy-number for all gene clusters.

The larger genome size of the Chinese and BC isolates reflect a higher proportion of repetitive elements, an enrichment that in itself explains the difference observed in genome size. Principal components analysis in figure 3 gives the proportion and composition of repetive elements between the single *N*.*neomacrospora* isolates. Each dot represents an isolate, the size of the dot indicates the proportion of the genome that have been identified as repetitive elements with ReapeatMasker. With the exception of the European and Quebec populations which fall together, the populations is clearly separated based on the composition of repetive elements. The different sized genome does not differ in the number of protein-coding genes, the number of genes with a signal peptide or the number of CAZymes. Significant differences between the two groups were found in the GC content and the degree of genome assembly. The combined European and Quebec group (EU-QC) have the highest GC content and most complete assemblies with a mean GC value of 53.8%, a L50 of 137 and mean number of contigs of 1,574, these values are contrasted by mean values for the British Columbia – China group (BC-CN): GC: 52.1%, L50: 172 and 6,483 contigs (Table 2). The gene count, number of CAZymes, and the number of genes with a signal peptide are comparable across the species, and no differentiation was found between populations.

**Figure 3.**
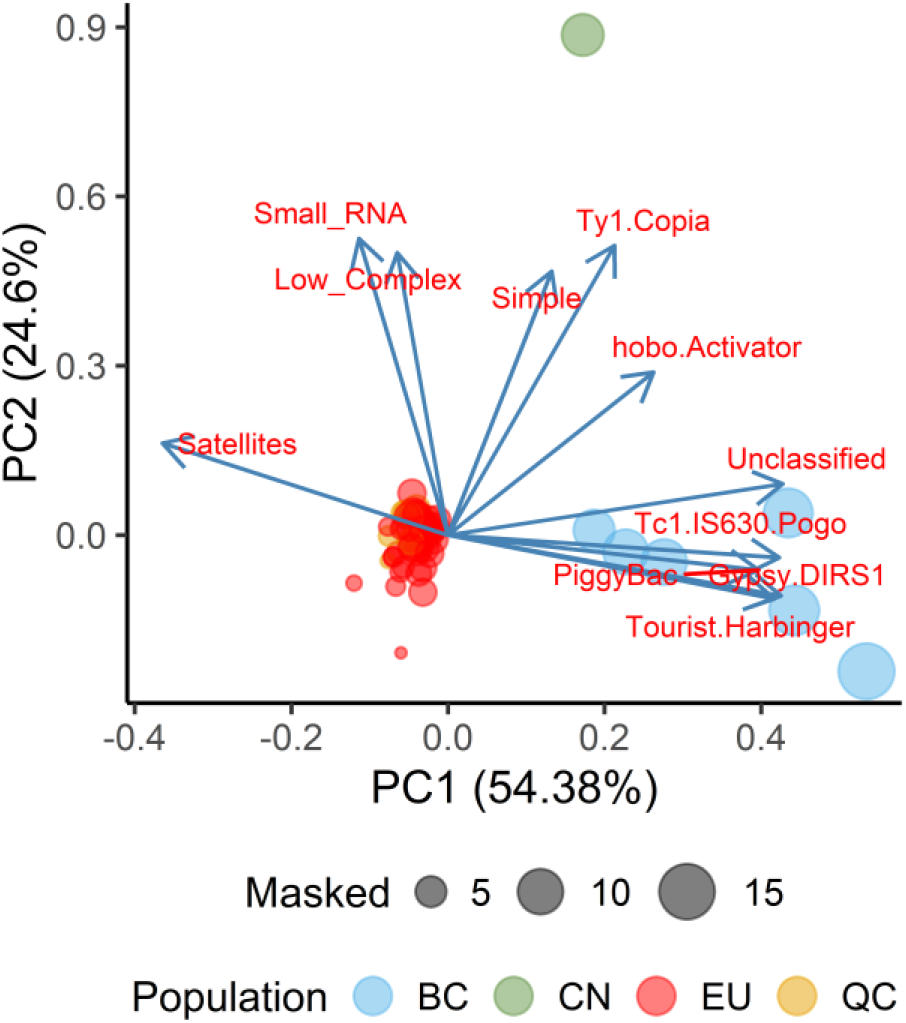
Principal components analysis on the proportion and composition of repetive elements between the single *Neonectria neomacrospora* isolates. Each dot represents an isolate, the size of the dot indicates the proportion of the genome that have been identified as repetitive elements with ReapeatMasker. The dot colours, show the geographical origin of the isolate, BC: British Columbia (n=6), CH: China (n=1), EU: Europe (n=38) and QC: Quebec (n=15). The Quebec isolates are below the European dots.

The maximum likelihood phylogeny of the Nectriaceae species and strains included in this study, based on 51 single-copy genes, showed that the BC-CN lineages are ancestral to the monophyletic clade with the EU-QC lineages. All shown splits have bootstrap values of 100/100 (Figure 4). The fact that the EU-QC group forms a monophyletic clade derived from the more ancestral clades of BC and China indicates that the differences in genome size are a product of genome reduction in the EU-QC clade. The genome size of the two most closely related species, *N. ditissima* and *N. major*, range from 43.5-45.8 Mb (n=3) and 41.6-42.5 Mb (n=2), respectively. The genome size of *N. ditissima* is based on NCBI GenBank data, and the estimates for *N. major* is from this study, both estimates are based on isolates from two continents.

**Figure 4.**
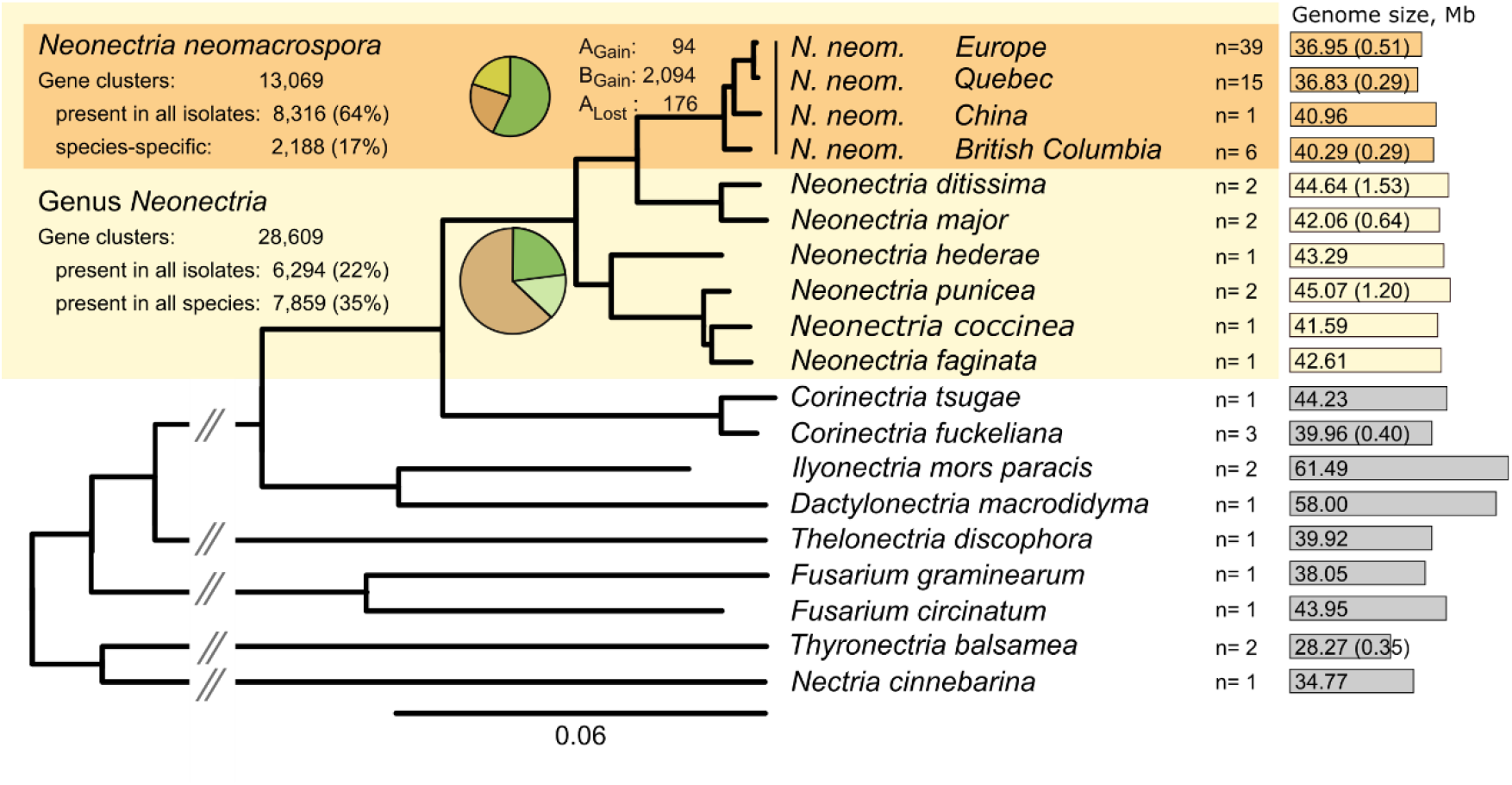
The maximum likelihood phylogeny of the 16 species within the family Nectriaceae. The phylogeny is based on 51 single-copy ortholog genes present in all strains. All splits shown have bootstrap values of 100/100. The computed phylogeny contained 83 strains, and the number of strains per species is noted to the right of the species name. The phylogeny is collapsed up to species level, except for N. neomacrospora four clades corresponding to the sampling region is maintained. Mean genome sizes and one standard deviation are given for each clade in the phylogeny. All genes identified were clustered based on their sequence similarity. The total number of gene clusters and the number of clusters shared by all isolates (core gene clusters) at genus and species level of *Neonectira neomacrospora* are annotated as well as their relative proportion of the total number of gene clusters. Core gene clusters at the genus level are given as clusters present in all strains sampled within the genus, and clusters in at least one strain per species. 2,188 (17 %) of the gene cluster identified in *N. neomacrospora* were species-specific. The species-specific clusters are partitioned into core and accessory clusters, marked as AGain. and BGain. respectively. Genes present in all *Neonetria* species except for *N. neomacrospora* (lost genes) are noted as ALost.

A mean genome size of 36.9 Mb makes the European and Quebec *N. neomacrospora* the species within the genera with the smallest recorded genome. The species *N. ditissima* and *N. punicea* show the largest recorded genomes within the genus, with sizes around 45-46 Mb.

The total number of protein-coding genes identified in each of the 61 genomes ranged from 10,720 to 11,392 (Figure S3). The predicted genes were cluster into tribes based on similarity, resulting in 13,069 gene clusters, of which 8,316 (64%) were shared among all analyzed genomes of *N. neomacrospora* (Table 2). We define this set of gene clusters as the core genome. A proportion of the gene clusters contain multi-copy genes, and the total gene count is, therefore, higher than the number of gene clusters. If the maximum gene copy number of any isolate is summarized across all 13,069 clusters, we observe 15,101 genes, 34% more genes than what is present in the gapless reference genome. The rarefaction curves for the pan-and core-genome of *N. neomacrospora* do not saturate within the 61 strains sampled, as shown in figure 5.

**Figure 5.**
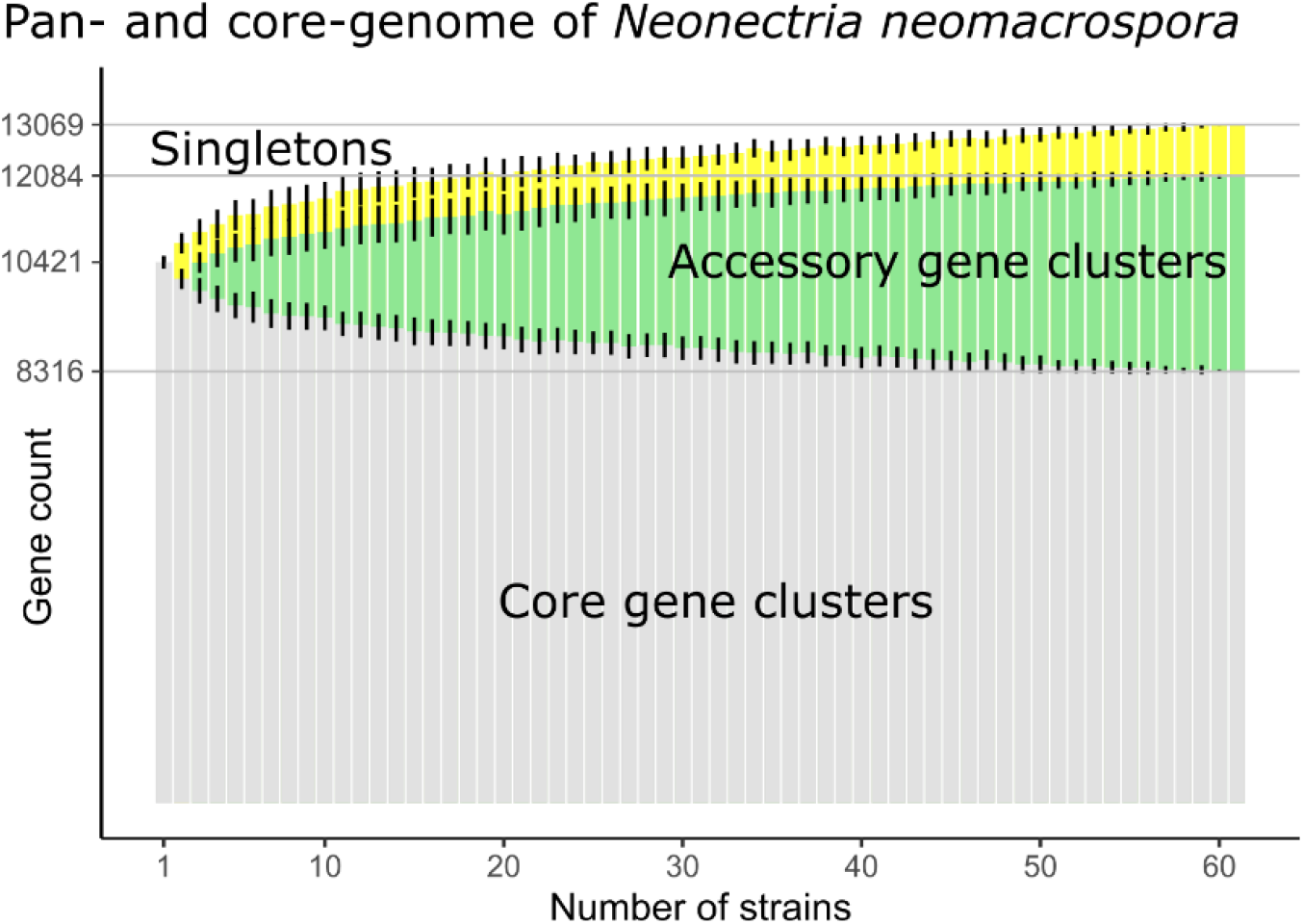
The pan-and core-genome of *Neonectria neomacrospora*. Estimation of the size of the core genome (as homolog gene clusters shared among all isolates) and the pangenome (all gene clusters). Gene clusters only found in one isolate (singletons) are shown in yellow. The total number of gene clusters identified in the 61 strains is 13,069, singletons composing 7.5% of these.

Of the total number of clusters within the pangenome, 2,188 (17%) are species-specific, 94 (0,7%) are present in all 61 sequenced *N. neomacrospora* genomes. One cluster had up to three gene copies per strain, and ten clusters had two gene copies per strain, equaling a total of 106 unique genes. The amount of species-specific gene clusters is similar in the two phylogenetic groups with 1,320 in the BC-CN group and 1,394 in the EU-QC group. 157 of the species-specific, and 15 of the species-specific core gene clusters contained gene with signal peptides. The reference genome contained 607 of the species-specific genes distributed across the genome. The density of species-specific genes was significantly higher in chr11 and chr12, compared to the larger chromosomes. Many of these genes have unknown functions and are likely effector candidates. The non-core part of the genome is with the exception of chr12, dispersed throughout the genome, not revealing an evident pattern. The core genome has a higher proportion of CAZymes and proteases compared to the non-core part. CAZymes: 4.5% vs 2.5% and proteases: 3.8% vs 2.8%. The proportion of protein sequences without any homologs in searched databases is slightly higher in the accessory part of the genome where 6.8 % of the protein sequences are unknown compared to 5.0% within the core part of the genome.

### CAZyme

We identified 457 putative genes that encode CAZymes in the genome of *N. neomacrospora*. Of the six CAZymes classes, carbohydrate esterases (CE), glycoside hydrolases (GH), and polysaccharide lyases (PL) are plant cell wall degrading enzymes (PCWDEs). There were 23, 221, and 16 genes belonging to the CE, GH, and PL classes, respectively. Three other classes with indirect roles in degrading carbohydrates are auxiliary activity (AA), carbohydrate-binding module (CBM), and glycosyl-transferase (GT). There were 58, 35, and 104 genes identified in the AA, CBM, and GT classes, respectively (Figure S4). CAZymes annotated to the strain KNNDK1 is distributed across all chromosomes, except chr12, with between 8.4 and 20.7 CAZymes (incl. AA and CBM) per Mb. (Table S4).

*Neonectria neomacrospora* have fewer PLs (range: 15-17) than the con-generic species (range: 21-26). Besides the two insect pathogenic fungi *Cordyceps militaris* and *Metarhizium anisopliae*, four other species have relative low PL counts: *Thyronectria balsamea* and *Corinectria fuckeliana, C. tsugae* and *Valsa mali*. Characteristic for the latter four is that they parasitize coniferous species, except *Valsa mali*. Fusarium circinatum is the conifer pathogen with the highest number of PLs with 18.

*Thelonectria discophora* was previously conceived as a cosmopolitical saprophyte but is now considered a complex of more restricted species (Salgado-Salazar *et al*., 2014). The isolate included in this study was isolated from an *Abies*, and it is possible that this strain also should be considered host limited to conifers. Its CAZyme profile, shown in figure 6, cluster it with *Corinectria* spp. and *N. neomacrospora*.

**Figure 6.**
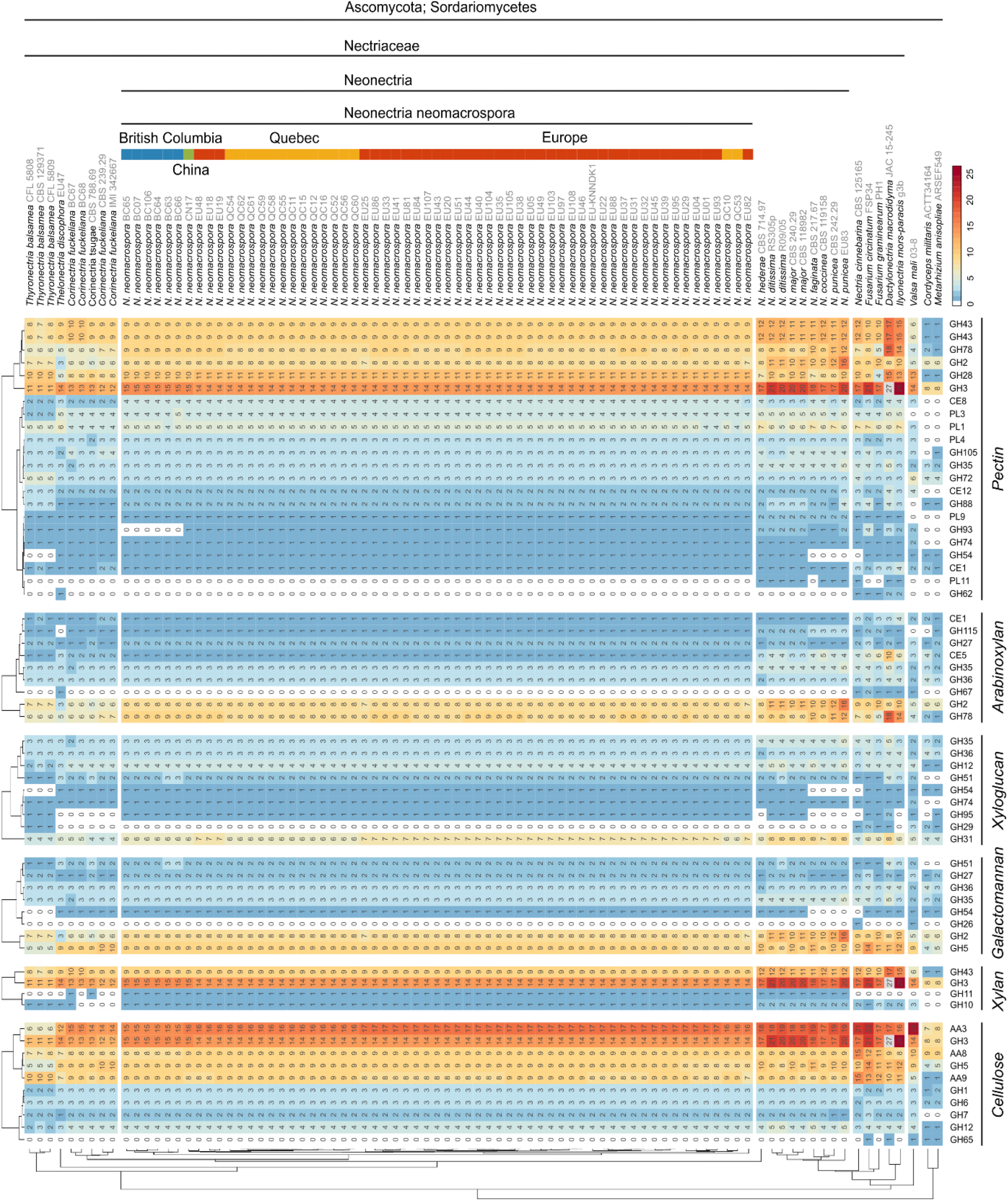
The CAZyme profile of *Neonectria neomacrospora* and related species within the Sordariomycetes. Strains are clustered hierarchically by their complete CAZyme profiles, including auxiliary activity molecules (AA) and carbohydrate-binding module (CBM) (not all data shown). A selection of enzymes and AA are presented according to the substrate within the plant cell wall they degrade: Cellulose, Pectin and the three hemicellulose compounds: Xylan, Xyloglucan and Arabinoxylan. Selection of enzyme families and assignment to substrate based on (Rytioja *et al*., 2014); included are carbohydrate esterases (CE), glycoside hydrolases (GH), polysaccharide lyases (PL) and auxiliary activity molecules (AA). Intra-species variation within *N. neomacrospora* is highlighted by the annotation of the geographical origin of each strain. *Thyronectria, Thelonectria, Corinectria, Fusarium circinatum* and *N. neomacrospora* strains are all isolated from coniferous hosts. The remaining *Neonectria* species are isolated from hardwood host species. *Cordyceps militaris* and *Metarhizium anisopliae* are insect pathogens.

The nine plant pathogenic fungal species analyzed that have hardwood or herbaceous crops as hosts have between 18 and 26 PLs (Figure 6). Analysis has not revealed enrichment of glucomannan specific enzymes in *N. neomacrospora* (glucomannan is a galactomannan), which could have been seen as a sign of adaptation to softwood degradation. The primary hemicellulose found in hardwood is not glucomannan as in softwood, but glucuronoxylan (Holtzapple, 2003), and while there is no enrichment of glucomannan specific enzymes, we see a depletion in both xylan enzymes (GH10, GH11, GH3, GH43) and the β-1,4-Galactosidase GH36, compared to the other species within the genus, all of which have hardwoods as host species. Cutinases are linked to pathogenicity in some species by facilitating the penetration of cuticle barriers in roots or leaves (Kolattukudy, 1985). The highest numbers of cutinases within the CE5 family observed in this study is also found in the root pathogens *Ilyonectria* and *Dactylonectria* species, and the leaf penetrating *Fusarium graminearum*, with between six and ten members of this enzyme family (Figure 6). *N. neomacrospora* has only a single CE5 enzyme; the lowest number observed and consistent with the perception of *N. neomacrospora* as a wound parasite.

The clustering of strains based on the abundance of the different GH perfectly separates species and *N. neomacrospora* into clades corresponding to geographical origin (data not shown), where the clustering based on all PCWDE, AA and CBM only separated BC form the remaining sample of *N. neomacrospora*. Four GH families showed consistent variation between the *N. neomacrospora* populations. The pectin degrading GH93 is present in all plant pathogens strains analysed with one to three enzymes (one in *N. neomacrospora*), but has seemingly been lost in the BC population of *N. neomacrospora*. The large GH3 family responsible for both cellulose and xylan degradation has been reduced from 15 to 14, in the monophyletic QC-EU group. GH28 includes ten enzymes in the CN and BC sample as well as in the sister species *N. ditissima* and *N. major*, but have expanded to eleven enzymes in the monophyletic group of QC and EU. The xyloglucan degrading GH31 family has six enzymes in all strains sampled from BC, CN and QC, but has seven across the European population.

The maximum distances between CAZyme profiles within *N. neomacrospora* (branch lengths of hclust-trees in figure 6) equal the distances observed between species such as *N. ditissima* and *N. major*, or *Corinectria fuckeliana* and *C. tsugae*.

## DISCUSSION

We assembled the genome of *N. neomacrospora* into twelve gapless chromosomes, comprising a total length of 37.1 Mb and 11,291 protein-coding genes. This places the genome size of *N. neomacrospora* close to the fungal median of 35 Mb, and 11,000 protein-coding genes, based on a sampling of 325 phylogenetically diverse fungi (Stajich, 2017). The mitochondrial genome of *N. neomacrospora* is approximately 22 Kb.

Telomere length between 107 bp and 142 bp, are within the expected 100-200 bp for filamentous fungi (Erlendson, Friedman and Freitag, 2017). The centromeres of most fungi are composed of a complex, heterogeneous set of repetitive sequences enriched in both AT-content and retrotransposons (Smith *et al*., 2012; Erlendson, Friedman and Freitag, 2017). Stretches with this pattern are present once in each of the assembled chromosomes. Centromeres can be identified based on these features in the representation of the 12 chromosomes in figure 1.

Centromere sequence structure is not well described for filamentous fungi, mainly due to its highly repetitive nature, which has impeded sequencing and assembly (Smith *et al*., 2012). With read-lengths, up to 60 kb in third-generation sequencing, assembling repetitive regions is not the obstacle it used to be. Recently, all seven centromeres of the rice blast fungus *Magnaporthe oryzae* were sequenced with PacBio. Highly AT-rich and heavily methylated DNA sequences were the only common defining features of the 57 kb to 109 kb *M. oryzae* centromere (Yadav *et al*., 2019). Within the Nectriaceae, several species in the well-described genus *Fusarium* have chromosomal regions that are not only AT-rich but also enriched in retrotransposons, and span 30-50 kb (Erlendson, Friedman and Freitag, 2017).

The observed differences between chromosomes fit the concept of a bipartite genome architecture with a non-random aggregation of genomic features. The apparent absence of chr12 in three out of the 61 isolates indicates that this chromosome is not essential, but can be considered as an accessory chromosome which can be either present or absent. The larger genome size observed in the strains sampled in British Columbia and China might be explained by the expansion of the repeat content. It is also possible that parts of the 953 gene clusters only found in the BC-CN group constitute a yet undescribed thirteenth chromosome.

Effectors are bioinformatically identified on a relatively broad criteria, which principally rely on the presence of a secretion signal, the N-terminal signal peptide motif, and the fact that the majority of effector proteins are small in size and rich in cysteine (Sperschneider *et al*., 2015). But not all secreted small cysteine-rich proteins will function as an effector, and not all effectors will fit this description and this approach is therefore problematic (Selin *et al*., 2016). More analysis is needed to determine whether the two minor chromosomes are enriched in effectors.

The four geographical defined populations split phylogenetically into four clades reflecting the geographic origin. The European and Quebec lineages form a monophyletic group, derived from the more ancestral lineages of China and British Columbia. It appears that the EU-QC clade has undergone a genome reduction, which might explain the historically reported phenotypic differences between BC and elsewhere. We have shown that there are substantial intra-species differences that could form a basis for dividing *Neonectria neomacrospora* into two races based on significant differences in genome size, repeat content, CAZyme combosition and deep phylogenetic split.

Ouellette (1972) found that isolates collected in Quebec caused more damage the infected host than isolates from BC and Europe. The present study includes some of the strains from Quebec and Norway used by Ouellette, as well as contemporary isolates collected from the same locations. None of the conducted analyses shaw a temporal structuring. We take this to mean that the 50 years past is negligible in this context and that conclusions from the Ouellette study can be interpreted in the light of our study. We found the Quebec, and European lineages to be mostly identical in terms of genomic architecture and found no discernible difference that could explain the phenotypic difference described by Ouellette.

It is noteworthy, that it is the BC race with the larger genomes that is considered avirulent. Some obligate biotrophic fungi have reduced genomes, whereas others show an increase in genome size primarily fueled by an increase in the effector repertoire to manipulate host metabolism and evade host immune response, facilitating the biotrophic lifestyle (N. Zhang *et al*., 2018). It could be speculated that a reduction in effector repertoire with the genome reduction has shifted the fungus from predominately biotrophic to predominately necrotrophic lifestyle, by shorting the latent phase where the fungus invades the host without causing symptoms. This is one possible explanation for the difference in virulence observed between BC, Quebec and Europe. There were, however, not observed a higher number of genes with a signal peptide in BC-CN genomes, compared to the reduced genomes of EU and Quebec.

The rapid evolution of effectors has been linked to TEs and it should be investigated to what extent the larger genome sizes of isolates for BC and China are driven by TEs. As illustrated in figure 1, TE rich regions of the genome have reduced GC content, something which is best explained by an active RIP silencing. It should be investigated, whether the lower CG content of the larger genomes from BC and China, is a sign of a higher proportion of TE. TE can drive genome size differences, as shown in *Zymoseptoria tritici* (Plissonneau, Hartmann and Croll, 2018). A possible signature of a high TE content lies in the significantly more fragmented *de novo* genome assemblies of the larger genomes. The BC-CN genomes are approximately 10 per cent larger than the smaller EU-QC genomes, but their fragmented assemblies consist of three times as many contigs. Repetitive regions are known to be difficult to assemble, and the observed fragmentation could, therefore, be explained by a relatively higher amount of TE. With the exception of the strain *N. hederae* CBS 714.97, then all species within the *Neonectria* genus are enriched with genes with predicted signal peptides compared to the twelve other species analysed (Figure S3). The analysed *N. neomacrospora* isolates have on average 978 (range: 915 to 1,019) genes with predicted signal peptides, three times as many as the insect pathogens *Cordyceps militaris* and *Metarhizium anisopliae*, and significantly more than the *Fusarium graminearum* and *F. circinatum* with 562 and 547, respectively. Future research will qualify candidates effector using programs like ApoplastP (Sperschneider *et al*., 2018) with throughout machine learning for prediction of effectors and their cellular localization. Knowledge on whether the candidate effectors are secreted cytoplastic or apoplastic can inform inferences on the biology of the species by comparative analysis.

Although *Neonectia* species usually are considered exclusively to be bark parasites, *N. coccinea* has been isolated from leaf litter of Beech (Unterseher, Peršoh and Schnittler, 2013). If present in the litter, were they then also there when the leaf was sitting on the tree? Questions like these shed light on how little we know about the life of these fungi. In working with this *N. neomacrospora*, we have numerous times observed perithecia of *N. neomacrospora* appear from the bark of newly felled trees that appeared healthy, or only lightly infected in a different part of the tree. It is possible that this pathogen can live for long periods as a biotrophic endophyte.

Castlebury, Rossman and Hyten (2005) found ground for splitting *C. cylindroides* found in British Columbia from *C. cylindroides* isolates from Europe into distinct species. This was based on sequencing data from three genes and five isolates: two from BC isolates and three from EU isolates (BC: CBS 118984, CBS 11898 EU: CBS 189.61, CBS 198.62, CBS 324.61). The genes were the translation elongation factor 1-alpha (EF1-a), RNA polymerase II second largest subunit (RPB2), and the β-tubulin gene.

The phylogenetic distance between the monophyletic group of the European and Quebec populations and the lineage found in British Columbia and China inferred from 51 random single-copy genes, is not as long as the distance observed between any pair of recognized species within the genus. This does not in itself mean that the two groups are not separate species, and the revealed consistent difference in genome size between the two groups adds to the picture of two distinct species. A key component of the plant pathogenic secretome is the carbohydrate-active enzymes, facilitating both infections in the appressorial phase and necrotrophy by degrading cell walls (O’Connell *et al*., 2012). We observed population-level differences in the CAZyme content, reflecting the described phylogenetic distance between populations.

We conclude that the common perceptions of the fungus as a concern in Europe and as a weak wound parasite of no concern in British Columbia might reflect the genetic differences. The population-level differences within *N. neomacrospora* are considerable. Given that the type species for N. neomacrospora is an isolate from British Columbia, it could be considered to erect the monophyletic group of the European and Quebec lineages into a subspecies or even a new species. We believe that further research into the genetic variation of this species is prudent before advancing this issue.

### URLS

Funannotate pipeline, https://funannotate.readthedocs.io/en/latest/index.html; ggpubr, https://rpkgs.datanovia.com/ggpubr/; RepeatMasker, http://www.repeatmasker.org/

## ACKNOWLEDGEMENT

We thank Dr Wen-Ying Zhuang (Chinese Academy of Sciences, Beijing) for providing an isolate of *N. neomacrospora* from China. Anne Uimari (Natural Resources Institute Finland, Luke) for collecting and providing samples from Finland. Halvor Solheim, Venche Talgø and Jan-Ole Skage for samples from Norway. Sophie Schmitz (Walloon Agricultural Research Centre) provided an isolate from Belgium.

We thank the Danish National High-Throughput DNA Sequencing Centre for its services. The Danish Christmas Tree Association supported fieldwork and sequencing that made this work possible.

## DATA AVAILABILITY STATEMENT

Raw reads and genomes assemblies of the 61 isolates described in this study are available the European Nucleotide Archive under the study accession number: PRJEB41540. The authors declare that all data of this study are available from the corresponding author upon reasonable request.

## Notes

### Competing Interest Statement

The authors have declared no competing interest.

## References

Almagro Armenteros, J. J. et al. (2019) ‘SignalP 5.0 improves signal peptide predictions using deep neural networks’, Nature Biotechnology. Springer US, 37(4), pp. 420–423. doi: 10.1038/s41587-019-0036-z.

Altschul, S. F. et al. (1990) ‘Basic local alignment search tool’, Journal of Molecular Biology, 215(3), pp. 403–410. doi: 10.1016/S0022-2836(05)80360-2.

Bao, W., Kojima, K. and Kohany, O. (2015) ‘Repbase Update, a database of repetitive elements in eukaryotic genomes’, Mobile DNA, 6(11). Available at: https://www.girinst.org/.

Bao, Z. and Eddy, S. R. (2002) ‘Automated de novo identification of repeat sequence families in sequenced genomes’, Genome Research. doi: 10.1101/gr.88502.

Bateman, A. et al. (2017) ‘UniProt: the universal protein knowledgebase’, Nucleic Acids Research. Oxford University Press, 45(D1), pp. D158–D169. doi: 10.1093/nar/gkw1099.

Benedict, M. N. et al. (2014) ‘ITEP: An integrated toolkit for exploration of microbial pan-genomes’, BMC Genomics, 15(1), p. 8. doi: 10.1186/1471-2164-15-8.

Besemer, J. and Borodovsky, M. (2005) ‘GeneMark: Web software for gene finding in prokaryotes, eukaryotes and viruses’, Nucleic Acids Research, 33(SUPPL. 2), pp. 451–454. doi: 10.1093/nar/gki487.

Boetzer, M. and Pirovano, W. (2014) ‘SSPACE-LongRead: Scaffolding bacterial draft genomes using long read sequence information’, BMC Bioinformatics, 15(1), pp. 1–9. doi: 10.1186/1471-2105-15-211.

Buchfink, B., Xie, C. and Huson, D. H. (2015) ‘Fast and sensitive protein alignment using DIAMOND’, Nature Methods, 12(1), pp. 59–60. doi: 10.1038/nmeth.3176.

Busk, P. K. et al. (2017) ‘Homology to peptide pattern for annotation of carbohydrate-active enzymes and prediction of function’, BMC Bioinformatics, 18(1), p. 214. doi: 10.1186/s12859-017-1625-9.

Chernomor, O., von Haeseler, A. and Minh, B. Q. (2016) ‘Terrace Aware Data Structure for Phylogenomic Inference from Supermatrices’, Systematic Biology, 65(6), pp. 997–1008. doi: 10.1093/sysbio/syw037.

Croll, D. and McDonald, B. A. (2012) ‘The accessory genome as a cradle for adaptive evolution in pathogens’, PLoS Pathogens, 8(4), pp. 8–10. doi: 10.1371/journal.ppat.1002608.

Dallery, J.-F. et al. (2017) ‘Gapless genome assembly of Colletotrichum higginsianum reveals chromosome structure and association of transposable elements with secondary metabolite gene clusters’, BMC Genomics, 18(1), p. 667. doi: 10.1186/s12864-017-4083-x.

Van Dam, P. et al. (2017) ‘A mobile pathogenicity chromosome in Fusarium oxysporum for infection of multiple cucurbit species’, Scientific Reports, 7(1), pp. 1–15. doi: 10.1038/s41598-017-07995-y.

Delmont, T. O. and Eren, A. M. (2018) ‘Linking pangenomes and metagenomes: the Prochlorococcus metapangenome’, PeerJ, 6, p. e4320. doi: 10.7717/peerj.4320.

Derbyshire, M. et al. (2017) ‘The complete genome sequence of the phytopathogenic fungus Sclerotinia sclerotiorum reveals insights into the genome architecture of broad host range pathogens’, Genome Biology and Evolution, 9(3), pp. 593–618. doi: 10.1093/gbe/evx030.

Dong, S., Raffaele, S. and Kamoun, S. (2015) ‘The two-speed genomes of filamentous pathogens: Waltz with plants’, Current Opinion in Genetics and Development. Elsevier Ltd, 35, pp. 57–65. doi: 10.1016/j.gde.2015.09.001.

Van Dongen, S. and Abreu-Goodger, C. (2012) ‘Using MCL to extract clusters from networks’, Methods in Molecular Biology. doi: 10.1007/978-1-61779-361-5_15.

Doolittle, W. F. and Zhaxybayeva, O. (2009) ‘On the origin of prokaryotic species’, Genome Research, 19(5), pp. 744–756. doi: 10.1101/gr.086645.108.

Eddy, S. R. (2011) ‘Accelerated Profile HMM Searches’, PLoS Computational Biology. edited by W. R. Pearson, 7(10), p. e1002195. doi: 10.1371/journal.pcbi.1002195.

Edgar, R. C. (2004) ‘MUSCLE: A multiple sequence alignment method with reduced time and space complexity’, BMC Bioinformatics, 5, p. 113. doi: 10.1186/1471-2105-5-113.

Ellinghaus, D., Kurtz, S. and Willhoeft, U. (2008) ‘LTRharvest, an efficient and flexible software for de novo detection of LTR retrotransposons’, BMC Bioinformatics, 9(1), p. 18. doi: 10.1186/1471-2105-9-18.

English, A. C. et al. (2012) ‘Mind the Gap: Upgrading Genomes with Pacific Biosciences RS Long-Read Sequencing Technology’, PLoS ONE, 7(11), pp. 1–12. doi: 10.1371/journal.pone.0047768.

EPPO (2017) Neonectria neomacrospora an emerging disease of fir trees in Northern Europe: addition to the EPPO Alert List, EPPO Reporting Service - Pest & Diseases. Available at: https://gd.eppo.int/reporting/article-6088.

Eren, A. M. et al. (2015) ‘Anvi’o: An advanced analysis and visualization platform for ‘omics data’, PeerJ. doi: 10.7717/peerj.1319.

Erlendson, A. A., Friedman, S. and Freitag, M. (2017) ‘A Matter of Scale and Dimensions: Chromatin of Chromosome Landmarks in the Fungi’, Microbiology Spectrum, 5(4), pp. 1–43. doi: 10.1128/microbiolspec.FUNK-0054-2017.

Faino, L. et al. (2016) ‘Transposons passively and actively contribute to evolution of the two-speed genome 1 of a fungal pathogen The Netherlands Running title: Genome evolution by transposable elements’, Genome Research, pp. 1091–1100. doi: 10.1101/gr.204974.116.Freely.

Fouché, S., Plissonneau, C. and Croll, D. (2018) ‘The birth and death of effectors in rapidly evolving filamentous pathogen genomes’, Current Opinion in Microbiology. doi: 10.1016/j.mib.2018.01.020.

Frantzeskakis, L. et al. (2018) ‘Signatures of host specialization and a recent transposable element burst in the dynamic one-speed genome of the fungal barley powdery mildew pathogen’, BMC Genomics. BMC Genomics, 19(1), pp. 1–23. doi: 10.1186/s12864-018-4750-6.

Gómez-Cortecero, A. et al. (2016) ‘Variation in Host and Pathogen in the Neonectria/Malus Interaction; toward an Understanding of the Genetic Basis of Resistance to European Canker’, Frontiers in Plant Science, 7(September), pp. 1–14. doi: 10.3389/fpls.2016.01365.

Haas, B. J. et al. (2008) ‘Automated eukaryotic gene structure annotation using EVidenceModeler and the Program to Assemble Spliced Alignments’, Genome Biology, 9(1), pp. 1–22. doi: 10.1186/gb-2008-9-1-r7.

Hao, Z. et al. (2019) ‘RIdeogram: drawing SVG graphics to visualize and map genome-wide data on the idiograms’, RIdeogram: drawing SVG graphics to visualize and map genome-wide data on the idiograms, pp. 1–11. doi: 10.7287/peerj.preprints.27928v1.

Hawksworth, F. G. (1996) Dwarf mistletoes: Biology, pathology and systematics. Agricutlur, Wiens, D. Agricutlur. Washington, DC: United States Department of Agriculture, Forest Service.

Hoff, K. J. et al. (2016) ‘BRAKER1: Unsupervised RNA-Seq-based genome annotation with GeneMark-ET and AUGUSTUS’, Bioinformatics, 32(5), pp. 767–769. doi: 10.1093/bioinformatics/btv661.

Holtzapple, M. T. (2003) ‘Hemicelluloses’, The Encyclopedia of Food Sciences and Nutrition. Second Edi. doi: 10.1016/B0-12-227055-X/00589-7.

Hubley, R. et al. (2016) ‘The Dfam database of repetitive DNA families’, 44(November 2015), pp. 81–89. doi: 10.1093/nar/gkv1272.

Huerta-Cepas, J. et al. (2017) ‘Fast genome-wide functional annotation through orthology assignment by eggNOG-mapper’, Molecular Biology and Evolution. doi: 10.1093/molbev/msx148.

Huerta-Cepas, J. et al. (2019) ‘eggNOG 5.0: a hierarchical, functionally and phylogenetically annotated orthology resource based on 5090 organisms and 2502 viruses’, Nucleic Acids Research, 47(D1), pp. D309–D314. doi: 10.1093/nar/gky1085.

Hunt, M. et al. (2015) ‘Circlator: Automated circularization of genome assemblies using long sequencing reads’, Genome Biology. Genome Biology, 16(1), pp. 1–10. doi: 10.1186/s13059-015-0849-0.

Irelan, J. T. and Selker, E. U. (1996) ‘Gene silencing in filamentous fungi?: RIP, NIIP and quelling’, J. Genet., 75(3), pp. 313–324.

Jones, P. et al. (2014) ‘InterProScan 5: Genome-scale protein function classification’, Bioinformatics, 30(9), pp. 1236–1240. doi: 10.1093/bioinformatics/btu031.

Kalyaanamoorthy, S. et al. (2017) ‘ModelFinder: Fast model selection for accurate phylogenetic estimates’, Nature Methods, 14(6), pp. 587–589. doi: 10.1038/nmeth.4285.

Katoh, K. and Standley, D. M. (2013) ‘MAFFT multiple sequence alignment software version 7: Improvements in performance and usability’, Molecular Biology and Evolution, 30(4), pp. 772–780. doi: 10.1093/molbev/mst010.

Kluyver, T. et al. (2016) Jupyter Notebooks—a publishing format for reproducible computational workflows, Positioning and Power in Academic Publishing: Players, Agents and Agendas. doi: 10.3233/978-1-61499-649-1-87.

Kolattukudy, P. E. (1985) ‘Enzymatic penetration of the plant cuticle by fungal pathogens’, Ann. Rev. Phytopathol., 23(49), pp. 223–50.

Kolde, R. (2018) ‘pheatmap: pretty heatmaps. R package version 1.0.10’. (CRAN). Koren, S. et al. (2017) ‘Canu: Scalable and accurate long-read assembly via adaptive κ-mer weighting and repeat separation’, Genome Research, 27(5), pp. 722–736. doi: 10.1101/gr.215087.116.

Li, H. et al. (2009) ‘The Sequence Alignment/Map format and SAMtools’, Bioinformatics, 25(16), pp. 2078–2079. doi: 10.1093/bioinformatics/btp352.

Li, H. (2018) ‘Minimap2: Pairwise alignment for nucleotide sequences’, Bioinformatics, 34(18), pp. 3094–3100. doi: 10.1093/bioinformatics/bty191.

Lombard, V. et al. (2014) ‘The carbohydrate-active enzymes database (CAZy) in 2013’, Nucleic Acids Research, 42(D1), pp. 490–495. doi: 10.1093/nar/gkt1178.

Lowe, T. M. and Eddy, S. R. (1997) ‘tRNAscan-SE: A Program for Improved Detection of Transfer RNA Genes in Genomic Sequence’, Nucleic Acids Research, 25(5), pp. 955–964. doi: 10.1093/nar/25.5.955.

McCarthy, C. G. P. and Fitzpatrick, D. A. (2019) ‘Pan-genome analyses of model fungal species’, Microbial Genomics, 5(2). doi: 10.1099/mgen.0.000243.

Mikheenko, A. et al. (2018) ‘Versatile genome assembly evaluation with QUAST-LG’, Bioinformatics, 34(13), pp. i142–i150. doi: 10.1093/bioinformatics/bty266.

Nguyen, L. T. et al. (2015) ‘IQ-TREE: A fast and effective stochastic algorithm for estimating maximum-likelihood phylogenies’, Molecular Biology and Evolution, 32(1), pp. 268–274. doi: 10.1093/molbev/msu300.

O’Connell, R. J. et al. (2012) ‘Lifestyle transitions in plant pathogenic Colletotrichum fungi deciphered by genome and transcriptome analyses.’, Nature genetics, 44(9), pp. 1060–5. doi: 10.1038/ng.2372.

Ouellette, G. B. (1972) ‘Nectria macrospora (Wr.) Ouellette sp. nov. (=N. fuckeliana var. macrospora): Strains, Physiology and Pathogenicity, and Comparison with N. fuckeliana var. fuckeliana’, Forest Pathology, 2(3), pp. 172–181. doi: 10.1111/j.1439-0329.1972.tb00358.x.

Pantou, M. P., Kouvelis, V. N. and Typas, M. A. (2008) ‘The complete mitochondrial genome of Fusarium oxysporum: Insights into fungal mitochondrial evolution’, Gene, 419(1–2), pp. 7–15. doi: 10.1016/j.gene.2008.04.009.

Plissonneau, C., Hartmann, F. E. and Croll, D. (2018) ‘Pangenome analyses of the wheat pathogen Zymoseptoria tritici reveal the structural basis of a highly plastic eukaryotic genome’, BMC Biology. BMC Biology, 16(1), pp. 1–16. doi: 10.1186/s12915-017-0457-4.

Porter, S. S. et al. (2017) ‘Association mapping reveals novel serpentine adaptation gene clusters in a population of symbiotic Mesorhizobium’, The ISME Journal. Nature Publishing Group, 11(1), pp. 248– 262. doi: 10.1038/ismej.2016.88.

Price, A. L., Jones, N. C. and Pevzner, P. A. (2005) ‘De novo identification of repeat families in large genomes’, Bioinformatics, 21(Suppl 1), pp. i351–i358. doi: 10.1093/bioinformatics/bti1018.

Raffaele, S. and Kamoun, S. (2012) ‘Genome evolution in filamentous plant pathogens: Why bigger can be better’, Nature Reviews Microbiology. Nature Publishing Group, 10(6), pp. 417–430. doi: 10.1038/nrmicro2790.

Rawlings, N. D., Barrett, A. J. and Finn, R. (2016) ‘Twenty years of the MEROPS database of proteolytic enzymes, their substrates and inhibitors’, Nucleic Acids Research, 44(D1), pp. D343–D350. doi: 10.1093/nar/gkv1118.

Reno, M. L. et al. (2009) ‘Biogeography of the Sulfolobus islandicus pan-genome’, Proceedings of the National Academy of Sciences, 106(21), pp. 8605–8610. doi: 10.1073/pnas.0808945106.

Rietman, L. M., Shamoun, S. F. and Van Der Kamp, B. J. (2005) ‘Forest pathology / Pathologie forestière Assessment of Neonectria neomacrospora (anamorph Cylindrocarpon cylindroides) as an inundative biocontrol agent against hemlock dwarf mistletoe’, Canadian Journal of Plant Pathology, 27, pp. 603– 609.

Robak, H. (1951) Noen iaktakelser til belysning av forholdet mellom klimatiske skader og soppangrep på nåletræer, Vestlandets Forstlige Forsøksstation. Bergen.

Rytioja, J. et al. (2014) ‘Plant-Polysaccharide-Degrading Enzymes from Basidiomycetes’, 78(4), pp. 614– 649. doi: 10.1128/MMBR.00035-14.

Salgado-Salazar, C. et al. (2014) ‘Phylogeny and taxonomic revision of Thelonectria discophora (Ascomycota, Hypocreales, Nectriaceae) species complex’, Fungal Diversity, 70(1), pp. 1–29. doi: 10.1007/s13225-014-0280-y.

Schubert, M., Lindgreen, S. and Orlando, L. (2016) ‘AdapterRemoval v2: Rapid adapter trimming, identification, and read merging’, BMC Research Notes. BioMed Central, 9(1), pp. 1–7. doi: 10.1186/s13104-016-1900-2.

Selin, C. et al. (2016) ‘Elucidating the role of effectors in plant-fungal interactions: Progress and challenges’, Frontiers in Microbiology, 7(APR), pp. 1–21. doi: 10.3389/fmicb.2016.00600.

Selker, E. U. et al. (1987) ‘Rearrangement of duplicated DNA in specialized cells of Neurospora’, Cell, 51(5), pp. 741–752. doi: 10.1016/0092-8674(87)90097-3.

Shen, W. et al./person-group>. (2016) ‘SeqKit: A Cross-Platform and Ultrafast Toolkit for FASTA/Q File Manipulation’, PLOS ONE. edited by Q. Zou, 11(10), p. e0163962. doi: 10.1371/journal.pone.0163962.

Simão, F. A. et al. (2015) ‘BUSCO: Assessing genome assembly and annotation completeness with single-copy orthologs’, Bioinformatics, 31(19), pp. 3210–3212. doi: 10.1093/bioinformatics/btv351.

Slater, G. S. C. and Birney, E. (2005) ‘Automated generation of heuristics for biological sequence comparison’, BMC Bioinformatics, 6, pp. 1–11. doi: 10.1186/1471-2105-6-31.

Smit, A., Hubley, R. and Green, P. (2015) ‘RepeatMasker’. Open-4.0. Available at: http://www.repeatmasker.org.

Smith, K. M. et al. (2012) ‘Centromeres of filamentous fungi’, Chromosome Research, 20(5), pp. 635– 656. doi: 10.1007/s10577-012-9290-3.

Spanu, P. D. et al. (2010) ‘Genome Expansion and Gene Loss in Powdery Mildew Fungi Reveal Tradeoffs in Extreme Parasitism’, Science, 330(6010), pp. 1543–1546. doi: 10.1126/science.1194573.

Sperschneider, J. et al. (2015) ‘Advances and Challenges in Computational Prediction of Effectors from Plant Pathogenic Fungi’, PLOS Pathogens, 11(5), p. e1004806. doi: 10.1371/journal.ppat.1004806.

Sperschneider, J. et al. (2018) ‘ApoplastP: prediction of effectors and plant proteins in the apoplast using machine learning’, New Phytologist, 217(4), pp. 1764–1778. doi: 10.1111/nph.14946.

Stajich, J. E. (2017) ‘Fungal Genomes and Insights into the Evolution of the Kingdom’, The Fungal Kingdom, pp. 619–633. doi: 10.1128/microbiolspec.funk-0055-2016.

Stanke, M. and Morgenstern, B. (2005) ‘AUGUSTUS?: a web server for gene prediction in eukaryotes that allows user-defined constraints’, 33, pp. 465–467. doi: 10.1093/nar/gki458.

Unterseher, M., Peršoh, D. and Schnittler, M. (2013) ‘Leaf-inhabiting endophytic fungi of European Beech (Fagus sylvatica L.) co-occur in leaf litter but are rare on decaying wood of the same host’, Fungal Diversity, 60(1), pp. 43–54. doi: 10.1007/s13225-013-0222-0.

Wickham, H. (2016) ggplot2: Elegant Graphics for Data Analysis. New York: Springer-Verlag. Available at: https://ggplot2.tidyverse.org.

Wollenweber, H. W. (1913) ‘Ramularia, Mycosphaerella, Nectria, Calorectria’, in Phytopathology. Baltimore, pp. 198–243.

Wu, C. et al. (2009) ‘Characterization of chromosome ends in the filamentous fungus Neurospora crassa’, Genetics, 181(3), pp. 1129–1145. doi: 10.1534/genetics.107.084392.

Yadav, V. et al. (2019) ‘Cellular dynamics and genomic identity of centromeres in cereal blast fungus’, mBio, 10(4), pp. 1–22. doi: 10.1128/mBio.01581-19.

Yin, Z. et al. (2015) ‘Genome sequence of Valsa canker pathogens uncovers a potential adaptation of colonization of woody bark’, New Phytologist, 208(4), pp. 1202–1216. doi: 10.1111/nph.13544.

Zhang, H. et al. (2018) ‘dbCAN2: a meta server for automated carbohydrate-active enzyme annotation’, Nucleic Acids Research, 46(W1), pp. W95–W101. doi: 10.1093/nar/gky418.

Zhang, N. et al. (2018) ‘Genome wide analysis of the transition to pathogenic lifestyles in Magnaporthales fungi’, Scientific Reports. Springer US, 8(1), pp. 1–13. doi: 10.1038/s41598-018-24301-6.

