## Supplementary material for "The pangenome of the fungal pathogen *Neonectria neomacrospora*"

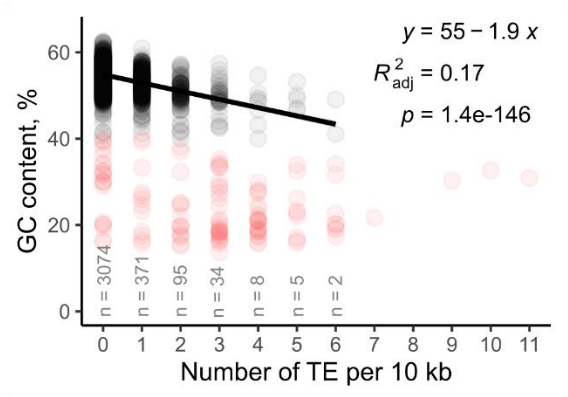

**Figure S1 | Guanine-cytosine (GC) content as a function of the number of transposable elements (TE).** 3706, 10 kb windows across the gemome of *N. neomacrospora* strain KNNDK1. Balck dots (n=3589) represents windows with a GC-content above 40%, the lower threshold for inclusion in the regression analysis. Red dots (n=117): GC-content below 40%, including centromeres and telomeres.

| Gene count | Signal peptides | |
|---|---|---|
| 11282 | 982 | *N. neomacrospora* BC106 |
| 10970 | 952 | *N. neomacrospora* BC63 |
| 11269 | 994 | *N. neomacrospora* BC64 |
| 11241 | 983 | *N. neomacrospora* BC65 |
| 11250 | 1005 | *N. neomacrospora* BC66 |
| 11267 | 988 | *N. neomacrospora* BC07 |
| 11357 | 1019 | *N. neomacrospora* CN17 |
| 10989 | 952 | *N. neomacrospora* EU01 |
| 11300 | 977 | *N. neomacrospora* EU103 |
| 11275 | 978 | *N. neomacrospora* EU104 |
| 11286 | 982 | *N. neomacrospora* EU105 |
| 11297 | 987 | *N. neomacrospora* EU107 |
| 11262 | 980 | *N. neomacrospora* EU108 |
| 10720 | 956 | *N. neomacrospora* EU18 |
| 10735 | 961 | *N. neomacrospora* EU19 |
| 11264 | 978 | *N. neomacrospora* EU20 |
| 11305 | 982 | *N. neomacrospora* EU22 |
| 11235 | 966 | *N. neomacrospora* EU25 |
| 11001 | 953 | *N. neomacrospora* EU29 |
| 11234 | 981 | *N. neomacrospora* EU31 |
| 11259 | 993 | *N. neomacrospora* EU32 |
| 11272 | 993 | *N. neomacrospora* EU33 |
| 11233 | 978 | *N. neomacrospora* EU35 |
| 11303 | 981 | *N. neomacrospora* EU37 |
| 11272 | 985 | *N. neomacrospora* EU38 |
| 11264 | 980 | *N. neomacrospora* EU39 |
| 11006 | 949 | *N. neomacrospora* EU04 |
| 11286 | 989 | *N. neomacrospora* EU40 |
| 11316 | 985 | *N. neomacrospora* EU41 |
| 11260 | 987 | *N. neomacrospora* EU43 |
| 11238 | 979 | *N. neomacrospora* EU44 |
| 11296 | 981 | *N. neomacrospora* EU45 |
| 11299 | 980 | *N. neomacrospora* EU46 |
| 11270 | 984 | *N. neomacrospora* EU48 |
| 11297 | 976 | *N. neomacrospora* EU49 |
| 11340 | 985 | *N. neomacrospora* EU05 |
| 11283 | 985 | *N. neomacrospora* EU51 |
| 11392 | 978 | *N. neomacrospora* EU81 |
| 10729 | 915 | *N. neomacrospora* EU82 |
| 11351 | 985 | *N. neomacrospora* EU84 |
| 11282 | 988 | *N. neomacrospora* EU86 |
| 11258 | 995 | *N. neomacrospora* EU88 |
| 11019 | 956 | *N. neomacrospora* EU93 |
| 11314 | 989 | *N. neomacrospora* EU95 |
| 11279 | 996 | *N. neomacrospora* EU97 |
| 11291 | 987 | *N. neomacrospora* EU-KNNDK1 |
| 10948 | 942 | *N. neomacrospora* QC10 |
| 11236 | 961 | *N. neomacrospora* QC11 |
| 11188 | 984 | *N. neomacrospora* QC12 |
| 11251 | 988 | *N. neomacrospora* QC15 |
| 11253 | 977 | *N. neomacrospora* QC16 |
| 11247 | 986 | *N. neomacrospora* QC52 |
| 10958 | 938 | *N. neomacrospora* QC53 |
| 11219 | 992 | *N. neomacrospora* QC54 |
| 11233 | 972 | *N. neomacrospora* QC55 |
| 11331 | 980 | *N. neomacrospora* QC56 |
| 11227 | 985 | *N. neomacrospora* QC58 |
| 11289 | 979 | *N. neomacrospora* QC59 |
| 11219 | 983 | *N. neomacrospora* QC60 |
| 11247 | 988 | *N. neomacrospora* QC61 |
| 11243 | 984 | *N. neomacrospora* QC62 |
| 12638 | 943 | *N. ditissima* RS305p |
| 12626 | 921 | *N. ditissima* R09/05 |
| 12894 | 971 | *N. major* CBS 240.29 |
| 12672 | 951 | *N. major* CBS 118982 |
| 11529 | 819 | *N. hederae* CBS 714.97 |
| 12645 | 982 | *N. punicea* CBS 242.29 |
| 13933 | 930 | *N. punicea* EU83 |
| 12534 | 904 | *N. coccinea* CBS 119158 |
| 12725 | 889 | *N. faginata* CBS 217.67 |
| 11508 | 716 | Corinectria tsugae CBS 788.69 |
| 11501 | 768 | *Corinectria fuckeliana* BC67 |
| 11459 | 765 | *Corinectria fuckeliana* BC68 |
| 11240 | 766 | *Corinectria fuckeliana* CBS 239.29 |
| 10972 | 731 | *Corinectria fuckeliana* IMI 342667 |
| 16006 | 723 | *Ilyonectria mors-paracis* g3b |
| 16255 | 813 | *Dactylonectria macrodidyma* JAC 15-245 |
| 12216 | 597 | *Thelonectria discophora* EU47 |
| 12267 | 562 | *Fusarium graminearum* PH1 |
| 14069 | 547 | *Fusarium circinatum* FSP34 |
| 9414 | 413 | *Thyronectria balsamea* CBS 129371 |
| 9441 | 413 | *Thyronectria balsamea* CFL 5808 |
| 9479 | 416 | *Thyronectria balsamea* CFL 5809 |
| 11027 | 586 | *Nectria cinnebarina* CBS 125165 |
| 9411 | 275 | *Cordyceps militaris* ACTT34164 |
| 11108 | 326 | *Metarhizium anisopliae* ARSEF549 |
| 10829 | 282 | *Valsa mali* 03-8 |

British Columbia, China, Europe, Quebec

*Neonectria neomacrospora*; *Neonectria*; Nectriaceae; Ascomycota; Sordariomycetes

**Figure S2 | List of the number of predicted genes and signal peptides with *Neonectria neomacrospora* and 18 other fungal species within the Sordariomycetes.** The 12 chromosomes of *Neonectria neomacrospora* strain KNNDK1 with pangenome annotation.

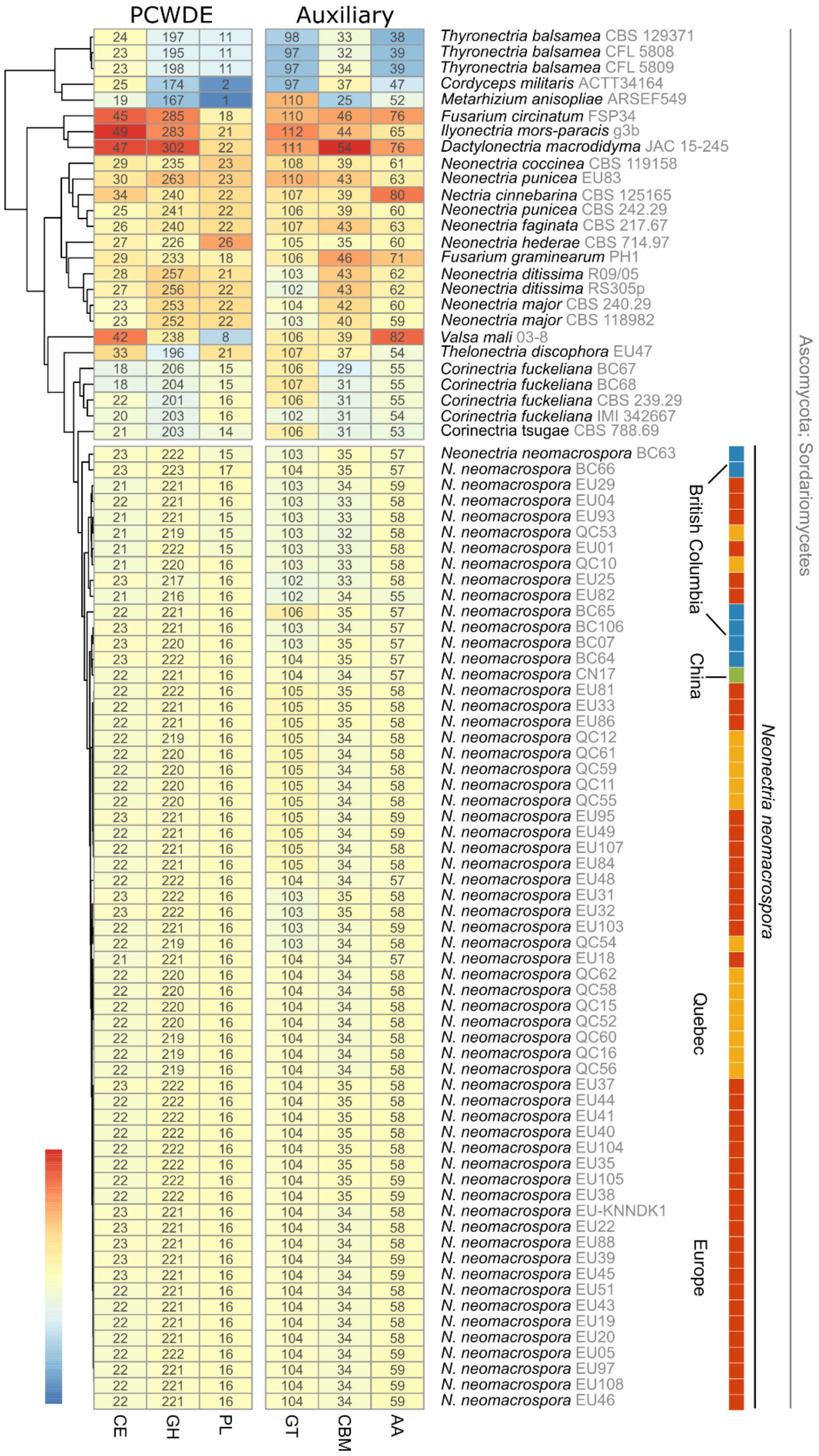

**Figure S3 | The CAZyme profile of *Neonectria neomacrospora* and related species within the Sordariomycetes.** Carbohydrate esterases (CE), glycoside hydrolases (GH), polysaccharide lyases (PL), and glycosyl-transferase (GT), carbohydrate-binding module (CBM) and auxiliary activity molecules (AA), is summarized per strain, and strains are clustered based on the summarized CAZyme profiles.

### Table S1 | Fungal isolates used in this study

| Species | * | Country | Location | Lat. | Long. | Year | Host | Source / Assesion no. | lab_ID | Collected or isolated by |
|---|---|---|---|---|---|---|---|---|---|---|
| *Neonectria neomacrospora* | EU | France | Vosges | 48.0707 | 6.9509 | 1957 | Abies alba | CBS 189.61 | 1 | W. Gerlach |
| *Neonectria neomacrospora* | EU | Netherlands | Zwolle | 52.5112 | 6.0940 | 1961 | Abies concolor | CBS 324.61 | 4 | L. Lombard |
| *Neonectria neomacrospora* | EU | Belgium | Herbeumont | 49.7689 | 5.2443 | 2017 | Abies grandis | BE5104 | 18 | S. Schmitz |
| *Neonectria neomacrospora* | EU | Switzerland | Arboretum | 46.5102 | 6.3684 | 2017 | Abies nordmaniana | CH01011 | 19 | K.N. Nielsen |
| *Neonectria neomacrospora* | EU | Norway | Os | 62.4965 | 11.2233 | 1958 | Abies alba | NO 1883/5 | 49 | Robak |
| *Neonectria neomacrospora* | EU | Norway | Fana | 60.2741 | 5.3954 | 1961 | Abies alba | CBS 503.67 | 5 | R. Roll-Hansen |
| *Neonectria neomacrospora* | EU | Norway | Fana | 60.2716 | 5.3866 | 1961 | Abies alba | NO 61-62/1 | 51 | R. Roll-Hansen |
| *Neonectria neomacrospora* | EU | Norway | Fana | 60.2600 | 5.3400 | 2019 | Abies lasiocarpa | NO 252125 | 93 | J.-O. Skage |
| *Neonectria neomacrospora* | EU | Norway | Fana | 60.2600 | 5.3400 | 2019 | Abies lasiocarpa | NO 252130 | 95 | J.-O. Skage |
| *Neonectria neomacrospora* | EU | Norway | Fana | 60.2600 | 5.3400 | 2019 | Abies lasiocarpa | NO 252140 | 97 | J.-O. Skage |
| *Neonectria neomacrospora* | EU | Denmark | Arboretum | 55.8691 | 12.5033 | 2015 | Abies fargesii | DK01011 | 20 | K.N. Nielsen |
| *Neonectria neomacrospora* | EU | Denmark | Arboretum | 55.8642 | 12.5119 | 2016 | Abies lasiocarpa | DK01081 | 22 | K.N. Nielsen |
| *Neonectria neomacrospora* | EU | Denmark | Arboretum | 55.8642 | 12.5093 | 2015 | Abies pinsapo | DK01132 | 25 | K.N. Nielsen |
| *Neonectria neomacrospora* | EU | Denmark | Silkeborg | 56.1634 | 9.5745 | 2015 | Abies nordmaniana | KNNDK1 | rf | K.N. Nielsen |
| *Neonectria neomacrospora* | EU | Denmark | Silkeborg | 56.1627 | 9.5750 | 2016 | Abies nordmaniana | DK02073 | 29 | K.N. Nielsen |
| *Neonectria neomacrospora* | EU | Denmark | Silkeborg | 56.1632 | 9.5757 | 2016 | Abies nordmaniana | DK02232 | 31 | K.N. Nielsen |
| *Neonectria neomacrospora* | EU | Denmark | Silkeborg | 56.1625 | 9.5745 | 2016 | Abies nordmaniana | DK02251 | 32 | K.N. Nielsen |
| *Neonectria neomacrospora* | EU | Denmark | Silkeborg | 56.1626 | 9.5741 | 2016 | Abies nordmaniana | DK02261 | 33 | K.N. Nielsen |
| *Neonectria neomacrospora* | EU | Denmark | Silkeborg | 56.1626 | 9.5718 | 2016 | Abies nordmaniana | DK02281 | 34 | K.N. Nielsen |
| *Neonectria neomacrospora* | EU | Denmark | Thy | 57.0242 | 8.5987 | 2015 | Abies nordmaniana | DK03011 | 37 | K.N. Nielsen |
| *Neonectria neomacrospora* | EU | Denmark | Thy | 57.0241 | 8.5989 | 2015 | Abies nordmaniana | DK03021 | 38 | K.N. Nielsen |
| *Neonectria neomacrospora* | EU | Denmark | Christiansfeld | 55.3643 | 9.4378 | 2018 | Abies procera | DK10021 | 39 | K.N. Nielsen |
| *Neonectria neomacrospora* | EU | Denmark | Christiansfeld | 55.3639 | 9.4378 | 2018 | Abies procera | DK10041 | 40 | K.N. Nielsen |
| *Neonectria neomacrospora* | EU | Denmark | Christiansfeld | 55.3637 | 9.4379 | 2018 | Abies procera | DK10051 | 41 | K.N. Nielsen |
| *Neonectria neomacrospora* | EU | Denmark | Christiansfeld | 55.3630 | 9.4378 | 2018 | Abies procera | DK10091 | 43 | K.N. Nielsen |
| *Neonectria neomacrospora* | EU | Denmark | Christiansfeld | 55.3626 | 9.4377 | 2018 | Abies procera | DK10101 | 44 | K.N. Nielsen |
| *Neonectria neomacrospora* | EU | Denmark | Christiansfeld | 55.3625 | 9.4378 | 2018 | Abies procera | DK10111 | 45 | K.N. Nielsen |
| *Neonectria neomacrospora* | EU | Denmark | Christiansfeld | 55.3621 | 9.4376 | 2018 | Abies procera | DK10121 | 46 | K.N. Nielsen |
| *Neonectria neomacrospora* | EU | Denmark | Bommerlund | 54.8790 | 9.3447 | 2018 | Abies nordmaniana | DK09011 | 81 | K.N. Nielsen |

| | | | | | | | | | | |
|---|---|---|---|---|---|---|---|---|---|---|
| *Neonectria neomacrospora* | EU | Denmark | Bommerlund | 54.8782 | 9.3442 | 2018 | Abies nordmaniana | DK09111 | 82 | K.N. Nielsen |
| *Neonectria neomacrospora* | EU | Denmark | Skelhusmarken | 56.7781 | 9.8417 | 2015 | Abies nordmaniana | DK04021 | 103 | K.N. Nielsen |
| *Neonectria neomacrospora* | EU | Denmark | Skelhusmarken | 56.7789 | 9.8423 | 2015 | Abies nordmaniana | DK04032 | 104 | K.N. Nielsen |
| *Neonectria neomacrospora* | EU | Denmark | Skelhusmarken | 56.7790 | 9.8420 | 2015 | Abies nordmaniana | DK04071 | 105 | K.N. Nielsen |
| *Neonectria neomacrospora* | EU | Denmark | Varde | 55.5957 | 8.5284 | 2016 | Abies grandis | DK07033 | 107 | K.N. Nielsen |
| *Neonectria neomacrospora* | EU | Denmark | Varde | 55.5880 | 8.5235 | 2016 | Abies grandis | DK07041 | 108 | K.N. Nielsen |
| *Neonectria neomacrospora* | EU | Finland | Mustila | 60.7315 | 26.4214 | 2018 | Abies sp. | FI01011 | 48 | A. Uimari |
| *Neonectria neomacrospora* | EU | Finland | Jarvenpaa | 60.4664 | 25.0896 | 2019 | Abies sp. | FI01021 | 84 | A. Uimari |
| *Neonectria neomacrospora* | EU | Finland | Espoo L 2 | 60.2014 | 24.8041 | 2019 | Abies sp. | FI01041 | 86 | A. Uimari |
| *Neonectria neomacrospora* | EU | Finland | Salo 15 | 60.3841 | 23.0868 | 2019 | Abies sp. | FI01061 | 88 | A. Uimari |
| *Neonectria neomacrospora* | QC | Canada | Anti-costi Island | 49.8200 | -64.3500 | 1967 | Abies balsamea | QFB19253/ CFL961 | 10 | P.O. Duguay |
| *Neonectria neomacrospora* | QC | Canada | Anti-costi Island | 49.8551 | -64.1037 | 1967 | Abies balsamea | QFB19255 / CFL962 | 11 | P.O. Duguay |
| *Neonectria neomacrospora* | QC | Canada | Anti-costi Island | 49.8551 | -64.1037 | 1967 | Abies balsamea | QFB19262 / CFL965 | 12 | P.O. Duguay |
| *Neonectria neomacrospora* | QC | Canada | Anti-costi Island | 49.7493 | -63.1044 | 2018 | Abies balsamea | CA01011 | 52 | |
| *Neonectria neomacrospora* | QC | Canada | Anti-costi Island | 49.8225 | -63.8326 | 2018 | Abies balsamea | CA01021 | 53 | |
| *Neonectria neomacrospora* | QC | Canada | Anti-costi Island | 49.7564 | -63.0759 | 2018 | Abies balsamea | CA01031 | 54 | K.N. Nielsen |
| *Neonectria neomacrospora* | QC | Canada | Anti-costi Island | 49.8266 | -63.8492 | 2018 | Abies balsamea | CA01041 | 55 | K.N. Nielsen |
| *Neonectria neomacrospora* | QC | Canada | Anti-costi Island | 49.5589 | -62.6887 | 2018 | Abies balsamea | CA01051 | 56 | K.N. Nielsen |
| *Neonectria neomacrospora* | QC | Canada | Anti-costi Island | 49.7563 | -63.0761 | 2018 | Abies balsamea | CA01061 | 58 | K.N. Nielsen |
| *Neonectria neomacrospora* | QC | Canada | Anti-costi Island | 49.8224 | -63.8323 | 2018 | Abies balsamea | CA01071 | 59 | K.N. Nielsen |
| *Neonectria neomacrospora* | QC | Canada | Anti-costi Island | 49.9146 | -64.3410 | 2018 | Abies balsamea | CA01081 | 60 | K.N. Nielsen |
| *Neonectria neomacrospora* | QC | Canada | Anti-costi Island | 49.9150 | -64.3394 | 2018 | Abies balsamea | CA01091 | 61 | K.N. Nielsen |
| *Neonectria neomacrospora* | QC | Canada | Port Cartier | 50.0300 | -66.8700 | 1967 | Abies balsamea | CFL 963 | 15 | |
| *Neonectria neomacrospora* | QC | Canada | Baie-Comeau | 49.2300 | -68.1600 | 1967 | Abies balsamea | CFL 964 | 16 | |
| *Neonectria neomacrospora* | QC | Canada | Cote Nord | 50.2998 | -66.4364 | 2018 | Abies balsamea | CA02011 | 62 | K.N. Nielsen |
| *Neonectria neomacrospora* | BC | Canada | British Columbia | 52.0000 | -126.0000 | 1996 | Tsuga heterophylla | CBS 118985 | 7 | S. Shamoun |
| *Neonectria neomacrospora* | BC | Canada | Cobble Hill | 48.7002 | -123.5912 | 2018 | Abies lasiocarpa | CA03011 | 63 | K.N. Nielsen |
| *Neonectria neomacrospora* | BC | Canada | Cobble Hill | 48.7005 | -123.5911 | 2018 | Abies lasiocarpa | CA03021 | 64 | K.N. Nielsen |
| *Neonectria neomacrospora* | BC | Canada | Cobble Hill | 48.7004 | -122.5910 | 2018 | Abies lasiocarpa | CA03031 | 65 | K.N. Nielsen |
| *Neonectria neomacrospora* | BC | Canada | Cobble Hill | 48.7002 | -123.5910 | 2018 | Abies lasiocarpa | CA03041 | 66 | K.N. Nielsen |
| *Neonectria neomacrospora* | BC | Canada | Vancouver Island | 49.3471 | -124.6259 | 2005 | Arceuthobium [illegible] | CBS 118984 | 106 | L. Reitman |

| | | | | | | | | | | |
|---|---|---|---|---|---|---|---|---|---|---|
| *Neonectria neomacrospora* | CN | China | Shennongjia | 31.7300 | 110.6800 | 2014 | Pinus sp. | HMAS 252906 | 17 | Zhao-Qing Zeng |
| *Neonectria ditissima* | | England | Kent | | | 2005 | Malus sp. | strain R09/05 \| GCA_001306435.1 | | |
| *Neonectria ditissima* | | New Zealand | Nelson | | | 2009 | Malus domestica | strain RS 305p \| GCA_001305495.1 | | |
| *Neonectria major* | | Norway | | 61.0000 | 8.5000 | | Alnus incana | CBS 240.29 | 3 | H.W. Wollenweber |
| *Neonectria major* | | USA | Washington | 46.6000 | -123.3000 | 2005 | Alnus rubra | CBS 118982 | 101 | C. Cootsma |
| *Neonectria coccinea* | | Germany | | 50.0000 | 10.0000 | | Fagus sylvatica | CBS 119158 | 8 | G.J. Samuels |
| *Neonectria faginata* | | Canada | New Brunswick | 46.5000 | -66.6000 | | Cryptococcus fagi nymph on | CBS 217.67 | 2 | G.L. Stone |
| *Neonectria punicea* | | Germany | | 50.0000 | 10.0000 | | Rhamnus sp. | CBS 242.29 | 100 | H.W. Wollenweber |
| *Neonectria punicea* | | Greece | | 37.0561 | 22.8124 | 2019 | Viscum album on Abies cephlonica | | 83 | M.J. Justesen |
| *Neonectria hederae* | | Netherlands | | | | 1932 | Hedera helix | CBS 714.97 \| GCA_003385265.1 | | J.W. Veenbaas-Rijks |
| *Corinectria fuckeliana* | | Canada | Joffre | 50.3516 | -122.4805 | 2018 | Abies sp. | | 67 | K.N. Nielsen |
| *Corinectria fuckeliana* | | Canada | Manning Park | 49.0591 | -120.8240 | 2018 | Abies sp. | | 68 | K.N. Nielsen |
| *Corinectria fuckeliana* | | Scotland | | 56.7000 | -4.3000 | | Picea sitchesis | CBS 239.29 | 69 | H.W. Wollenweber |
| *Corinectria fuckeliana* | | Switzerland | | | | 1990 | Unknown | IMI 342667 \| GCA_003385255.1 | | |
| *Corinectria tsugae* | | Canada | | 53.0000 | -96.0000 | 1966 | Tsuga heterophylla | CBS 788.69 | 70 | J.E. Bier |
| *Fusarium circinatum* | | USA | Califonia | | | 1996 | Pinus radiata | strain FSP 34 \| GCA_000497325.3 | | |
| *Fusarium graminearum* | | | | | | | | strain PH-1 \| GCA_900044135.1 | | |
| *Ilyonectria mors-paracis* | | | | | | | | strain g3b \| GCA_002991585.1 | | |
| *Dactylonectria macrodidyma* | | | | | | 2014 | Unknown | strain JAC 15-245 \| GCA_000935225.1 | | |
| *Thelonectria discophora* | | Denmark | Nødebo | 55.9884 | 12.3408 | 2017 | Abies procera | | 47 | K.N. Nielsen |
| *Thyronectria balsamea* | | Canada | Stittsville | 45.1996 | -75.9798 | 2009 | Abies balsamea | CBS 129371 | 102 | K.A. Seifert |
| *Thyronectria balsamea* | | Canada | La Patrie | 45.4026 | -71.2497 | 2016 | Abies balsamea | CFL 5809 | 14 | |
| *Thyronectria balsamea* | | Canada | Saint-Cyprien | 46.3308 | -70.2977 | 2018 | Abies balsamea | CFL 5808 | 13 | |
| *Nectria cinnabarina* | | France | Villiers en Bois | 46.1300 | -0.4100 | | Aesculus sp. | CBS 125165 | 74 | C. LeChat |
| *Metarhizium anisopliae* | | Brazil | | | | 1980 | Insect | strain ARSEF 549 \| GCA_000814975.1 | | |
| *Cordyceps militaris* | | | | | | | Insect Butterfly pupa | Strain ATCC 34164 \| GCA_008080495.1 | | |
| *Valsa mali* | | China | Shaanxi | | | 2006 | Malus sp. | strain 03-8 \| GCA_000818155.1 | | |

* Regional identifier, used to group isolates in the article

### Table S2 | Assembly statistics and read coverage

| Species | Country | Province | Collected | Source / Assesion no. | lab_ID | Coverage | #contigs | length | GC | N50 | N75 | L50 |
|---|---|---|---|---|---|---|---|---|---|---|---|---|
| *Neonectria neomacrospora* | France | Lac de Longmer, Vosges | 1957 | CBS 189.61 | 1 | 54.9 | 793 | 37021198 | 53.64 | 225535 | 135552 | 48 |
| *Neonectria neomacrospora* | Netherlands | Zwolle | 1961 | CBS 324.61 | 4 | 25.7 | 1015 | 37091102 | 53.65 | 236725 | 119474 | 53 |
| *Neonectria neomacrospora* | Belgium | Herbeumont | 2017 | BE5104 | 18 | 20.2 | 1917 | 36461970 | 54.28 | 65786 | 35561 | 170 |
| *Neonectria neomacrospora* | Switzerland | Jura | 2017 | CH01011 | 19 | 20.6 | 895 | 37151449 | 53.51 | 175620 | 90111 | 65 |
| *Neonectria neomacrospora* | Norway | Hordaland | 1958 | NO 1883/5 | 49 | 25.1 | 2047 | 36312304 | 54.41 | 54153 | 29584 | 197 |
| *Neonectria neomacrospora* | Norway | Hordaland | 1961 | CBS 503.67 | 5 | 27.0 | 887 | 37185101 | 53.59 | 199846 | 100159 | 56 |
| *Neonectria neomacrospora* | Norway | Hordaland | 1961 | NO 61-62/1 | 51 | 27.1 | 1588 | 36649618 | 54.23 | 131008 | 68412 | 90 |
| *Neonectria neomacrospora* | Norway | Hordaland | 2019 | NO 252125 | 93 | 25.0 | 1448 | 36953685 | 53.93 | 160193 | 85517 | 70 |
| *Neonectria neomacrospora* | Norway | Hordaland | 2019 | NO 252130 | 95 | 27.1 | 1693 | 36615896 | 54.21 | 116187 | 59460 | 95 |
| *Neonectria neomacrospora* | Norway | Hordaland | 2019 | NO 252140 | 97 | 24.0 | 1701 | 36571452 | 54.15 | 130597 | 67827 | 87 |
| *Neonectria neomacrospora* | Denmark | East Zealand | 2015 | DK01011 | 20 | 34.5 | 1223 | 36975782 | 53.79 | 192378 | 90265 | 64 |
| *Neonectria neomacrospora* | Denmark | East Zealand | 2016 | DK01081 | 22 | 19.4 | 1220 | 37053232 | 53.77 | 205333 | 101824 | 58 |
| *Neonectria neomacrospora* | Denmark | East Zealand | 2015 | DK01132 | 25 | 22.0 | 8714 | 38228153 | 53.77 | 117120 | 57303 | 93 |
| *Neonectria neomacrospora* | Denmark | East Jutland | 2016 | DK02073 | 29 | 38.5 | 663 | 37142615 | 53.56 | 291696 | 156980 | 42 |
| *Neonectria neomacrospora* | Denmark | East Jutland | 2016 | DK02232 | 31 | 31.1 | 1094 | 36934545 | 53.74 | 208103 | 117157 | 55 |
| *Neonectria neomacrospora* | Denmark | East Jutland | 2016 | DK02251 | 32 | 31.2 | 926 | 37076846 | 53.66 | 240590 | 135011 | 45 |
| *Neonectria neomacrospora* | Denmark | East Jutland | 2016 | DK02261 | 33 | 36.1 | 1854 | 37325201 | 53.61 | 252208 | 146648 | 47 |
| *Neonectria neomacrospora* | Denmark | East Jutland | 2016 | DK02281 | 35 | 29.0 | 1403 | 36943622 | 53.83 | 211433 | 128047 | 50 |
| *Neonectria neomacrospora* | Denmark | North Jutland | 2015 | DK03011 | 37 | 27.3 | 1552 | 36979382 | 53.82 | 177687 | 93370 | 62 |
| *Neonectria neomacrospora* | Denmark | North Jutland | 2015 | DK03021 | 38 | 27.4 | 1102 | 37173688 | 53.59 | 240501 | 120041 | 46 |
| *Neonectria neomacrospora* | Denmark | Southern Jutland | 2018 | DK10021 | 39 | 27.9 | 918 | 37095600 | 53.53 | 242851 | 139445 | 47 |
| *Neonectria neomacrospora* | Denmark | Southern Jutland | 2018 | DK10041 | 40 | 35.9 | 705 | 37114797 | 53.58 | 298831 | 152924 | 39 |
| *Neonectria neomacrospora* | Denmark | Southern Jutland | 2018 | DK10051 | 41 | 31.0 | 680 | 37269791 | 53.41 | 289122 | 161549 | 41 |
| *Neonectria neomacrospora* | Denmark | Southern Jutland | 2018 | DK10091 | 43 | 19.9 | 1038 | 37075113 | 53.61 | 201219 | 104527 | 53 |
| *Neonectria neomacrospora* | Denmark | Southern Jutland | 2018 | DK10101 | 44 | 27.6 | 1029 | 37100765 | 53.59 | 235967 | 137390 | 48 |

| | | | | | | | | | | | | |
|---|---|---|---|---|---|---|---|---|---|---|---|---|
| *Neonectria neomacrospora* | Denmark | Southern Jutland | 2018 | DK10111 | 45 | 25.3 | 713 | 37052615 | 53.59 | 225935 | 122065 | 48 |
| *Neonectria neomacrospora* | Denmark | Southern Jutland | 2018 | DK10121 | 46 | 22.5 | 977 | 37146474 | 53.56 | 216117 | 120842 | 56 |
| *Neonectria neomacrospora* | Denmark | Southern Jutland | 2018 | DK09011 | 81 | 16.5 | 2603 | 36488173 | 54.25 | 45725 | 24190 | 248 |
| *Neonectria neomacrospora* | Denmark | Southern Jutland | 2018 | DK09111 | 82 | 12.4 | 7523 | 36022668 | 54.44 | 8694 | 5077 | 1277 |
| *Neonectria neomacrospora* | Denmark | East Jutland | 2015 | DK04021 | 103 | 35.4 | 701 | 37160680 | 53.50 | 268646 | 164683 | 46 |
| *Neonectria neomacrospora* | Denmark | East Jutland | 2015 | DK04032 | 104 | 34.2 | 1013 | 37161798 | 53.66 | 263382 | 142176 | 45 |
| *Neonectria neomacrospora* | Denmark | East Jutland | 2015 | DK04071 | 105 | 32.7 | 978 | 37110878 | 53.58 | 225360 | 115094 | 51 |
| *Neonectria neomacrospora* | Denmark | West Jutland | 2016 | DK07033 | 107 | 19.8 | 1794 | 36488527 | 54.25 | 106266 | 56441 | 108 |
| *Neonectria neomacrospora* | Denmark | West Jutland | 2016 | DK07041 | 108 | 35.4 | 541 | 37062301 | 53.60 | 261055 | 142735 | 47 |
| *Neonectria neomacrospora* | Finland | Southern Finland | 2018 | FI01011 | 48 | 19.5 | 1075 | 37203405 | 53.58 | 220690 | 115344 | 52 |
| *Neonectria neomacrospora* | Finland | Southern Finland | 2019 | FI01021 | 84 | 28.7 | 1762 | 36452860 | 54.34 | 78007 | 43594 | 146 |
| *Neonectria neomacrospora* | Finland | Southern Finland | 2019 | FI01041 | 86 | 26.2 | 1841 | 36464338 | 54.26 | 78890 | 43080 | 137 |
| *Neonectria neomacrospora* | Finland | Southern Finland | 2019 | FI01061 | 88 | 21.5 | 998 | 37166452 | 53.57 | 184820 | 94892 | 61 |
| *Neonectria neomacrospora* | Canada | Quebec | 1967 | QFB19253 / CFL961 | 10 | 60.6 | 1185 | 36695899 | 53.76 | 191600 | 93673 | 56 |
| *Neonectria neomacrospora* | Canada | Quebec | 1967 | QFB19255 / CFL962 | 11 | 41.3 | 1416 | 36870701 | 53.78 | 143783 | 80213 | 74 |
| *Neonectria neomacrospora* | Canada | Quebec | 1967 | QFB19262 / CFL965 | 12 | 49.8 | 1391 | 36862386 | 53.71 | 179801 | 90456 | 61 |
| *Neonectria neomacrospora* | Canada | Quebec | 2018 | CA01011 | 52 | 27.7 | 1462 | 36679750 | 53.98 | 149872 | 79672 | 74 |
| *Neonectria neomacrospora* | Canada | Quebec | 2018 | CA01021 | 53 | 25.5 | 1685 | 37291616 | 53.54 | 188252 | 99432 | 55 |
| *Neonectria neomacrospora* | Canada | Quebec | 2018 | CA01031 | 54 | 30.5 | 864 | 37036564 | 53.56 | 223232 | 114296 | 51 |
| *Neonectria neomacrospora* | Canada | Quebec | 2018 | CA01041 | 55 | 83.0 | 972 | 36425792 | 54.20 | 142821 | 84433 | 79 |
| *Neonectria neomacrospora* | Canada | Quebec | 2018 | CA01051 | 56 | 32.7 | 1461 | 36964005 | 53.61 | 98032 | 52487 | 115 |
| *Neonectria neomacrospora* | Canada | Quebec | 2018 | CA01061 | 58 | 39.2 | 869 | 36953958 | 53.61 | 271815 | 132335 | 44 |
| *Neonectria neomacrospora* | Canada | Quebec | 2018 | CA01071 | 59 | 16.4 | 1910 | 36263659 | 54.40 | 58786 | 31512 | 192 |
| *Neonectria neomacrospora* | Canada | Quebec | 2018 | CA01081 | 60 | 24.3 | 985 | 37018946 | 53.61 | 188467 | 106612 | 57 |
| *Neonectria neomacrospora* | Canada | Quebec | 2018 | CA01091 | 61 | 18.7 | 1639 | 36370405 | 54.31 | 97585 | 52049 | 112 |
| *Neonectria neomacrospora* | Canada | Quebec | 1967 | CFL 963 | 15 | 25.5 | 850 | 37098179 | 53.48 | 224687 | 107262 | 56 |
| *Neonectria neomacrospora* | Canada | Quebec | 1967 | CFL 964 | 16 | 20.3 | 1387 | 36969610 | 53.60 | 118973 | 63750 | 93 |

| | | | | | | | | | | | | |
|---|---|---|---|---|---|---|---|---|---|---|---|---|
| *Neonectria neomacrospora* | Canada | Quebec | 2018 | CA02011 | 62 | 25.2 | 1605 | 36893067 | 53.82 | 158930 | 87451 | 65 |
| *Neonectria neomacrospora* | Canada | British Columbia | 1996 | CBS 118985 | 7 | 22.3 | 9490 | 42242818 | 50.91 | 76109 | 36528 | 165 |
| *Neonectria neomacrospora* | Canada | British Columbia | 2018 | CA03011 | 63 | 23.1 | 6576 | 39112794 | 52.94 | 64377 | 32101 | 177 |
| *Neonectria neomacrospora* | Canada | British Columbia | 2018 | CA03021 | 64 | 20.8 | 6175 | 38525360 | 53.41 | 50293 | 27196 | 219 |
| *Neonectria neomacrospora* | Canada | British Columbia | 2018 | CA03031 | 65 | 23.0 | 8552 | 40210002 | 52.41 | 56026 | 27875 | 205 |
| *Neonectria neomacrospora* | Canada | British Columbia | 2018 | CA03041 | 66 | 20.4 | 6543 | 39312554 | 52.77 | 61360 | 31090 | 184 |
| *Neonectria neomacrospora* | Canada | British Columbia | 2005 | CBS 118984 | 106 | 89.8 | 2558 | 40869945 | 51.18 | 98013 | 47611 | 125 |
| *Neonectria neomacrospora* | China | Shennongjia | 2014 | HMAS 252906 | 17 | 18.7 | 5484 | 40961935 | 51.39 | 90957 | 48970 | 130 |
| *Neonectria major* | Norway | | | CBS 240.29 | 3 | 17.3 | 5553 | 41611567 | 53.53 | 89485 | 49459 | 142 |
| *Neonectria major* | USA | Washington | 2005 | CBS 118982 | 101 | 21.4 | 11956 | 42516960 | 53.40 | 122776 | 65949 | 102 |
| *Neonectria coccinea* | Germany | | | CBS 119158 | 8 | 16.2 | 4031 | 41585127 | 52.76 | 129994 | 74995 | 103 |
| *Neonectria faginata* | Canada | New Brunswick | | CBS 217.67 | 2 | 15,9 | 4129 | 42605268 | 52.66 | 126099 | 64437 | 105 |
| *Neonectria punicea* | Germany | | | CBS 242.29 | 100 | 22.7 | 18205 | 45921251 | 51.25 | 121164 | 62734 | 108 |
| *Neonectria punicea* | Greece | | 2019 | | 83 | 19.5 | 5179 | 44222629 | 53.49 | 62750 | 33618 | 217 |
| *Corinectria fuckeliana* | Canada | British Columbia | 2018 | | 67 | 22.7 | 6624 | 39781634 | 53.16 | 71895 | 37001 | 162 |
| *Corinectria fuckeliana* | Canada | British Columbia | 2018 | | 68 | 18.9 | 8634 | 40424722 | 52.51 | 88016 | 45278 | 134 |
| *Corinectria fuckeliana* | Scotland | | | CBS 239.29 | 99 | 25.6 | 4622 | 39681832 | 52.35 | 65626 | 36250 | 177 |
| *Corinectria tsugae* | Canada | | 1966 | CBS 788.69 | 6 | | 19832 | 44227555 | 51.92 | 25567 | 11577 | 468 |
| *Thyronectria balsamea* | Canada | Ontario | 2009 | CBS 129371 | 102 | | 2572 | 28318405 | 51.81 | 100147 | 60880 | 85 |
| *Thyronectria balsamea* | Canada | Quebec | 2018 | CFL 5808 | 13 | | 3334 | 28815679 | 51.08 | 120563 | 62091 | 75 |
| *Nectria cinnabarina* | France | Villiers en Bois | | CBS 125165 | 9 | | 1201 | 34774977 | 51.32 | 169264 | 90735 | 61 |

Table S3 | Annotation for the 51 single-copy genes used for phylogenetic inference

| Chr | Start | End | | Product | Ontology_term | InterProScan and PFAM reference | note |
|---|---|---|---|---|---|---|---|
| chr3 | 4144044 | 4145497 | + | **26S proteasome regulatory subunit 6B** | GO_component: GO:0005737 - cytoplasm [Evidence IEA],GO_function: GO:0016787 - hydrolase activity [Evidence IEA],GO_function: GO:0005524 - ATP binding [Evidence IEA],GO_process: GO:0030163 - protein catabolic process [Evidence IEA] | InterPro:IPR003593,InterPro:IPR003959,InterPro:IPR003960,InterPro:IPR005937,InterPro:IPR027417,InterPro:IPR032501 PFAM:PF00004,PFAM:PF07724,PFAM:PF07728,PFAM:PF16450 | note=EggNog:ENOG410PGYA,COG:O |
| chr2 | 2432671 | 2434064 | - | **26S proteasome regulatory subunit rpn6** | | InterPro:IPR000717,InterPro:IPR036390,InterPro:IPR040773,InterPro:IPR040780,PFAM:PF01399,PFAM:PF18055,PFAM:PF18503 | note=BUSCO:EOG092630YS,EggNog:ENOG410PJ50,COG:O |
| chr2 | 2832680 | 2833181 | - | **40S ribosomal protein S20** | GO_component: GO:0005840 - ribosome [Evidence IEA],GO_component: GO:0015935 - small ribosomal subunit [Evidence IEA],GO_function: GO:0003735 - structural constituent of ribosome [Evidence IEA],GO_function: GO:0003723 - RNA binding [Evidence IEA],GO_process: GO:0006412 - translation [Evidence IEA] | InterPro:IPR001848,InterPro:IPR005729,InterPro:IPR018268,InterPro:IPR027486,InterPro:IPR036838,PFAM:PF00338 | note=BUSCO:EOG09265CFP,EggNog:ENOG410PQ4U,COG:J |
| chr1 | 2534170 | 2534807 | - | **60S ribosomal protein L11** | GO_component: GO:0005840 - ribosome [Evidence IEA],GO_function: GO:0003735 - structural constituent of ribosome [Evidence IEA],GO_process: GO:0006412 - translation [Evidence IEA] | InterPro:IPR002132,InterPro:IPR020929,InterPro:IPR022803,InterPro:IPR031309,InterPro:IPR031310 PFAM:PF00281,PFAM:PF00673 | note=BUSCO:EOG09264X8J,EggNog:ENOG410PGC7,COG:J |
| chr2 | 3997720 | 3998500 | - | **60S ribosomal protein L25** | GO_component: GO:0005840 - ribosome [Evidence IEA],GO_function: GO:0003735 - structural constituent of ribosome [Evidence IEA],GO_process: GO:0006412 - translation [Evidence IEA] | InterPro:IPR005633,InterPro:IPR012677,InterPro:IPR012678,InterPro:IPR013025,PFAM:PF00276,PFAM:PF03939 | note=BUSCO:EOG092657YR,EggNog:ENOG410PPAS,COG:J |
| chr2 | 4488547 | 4489250 | - | **60S ribosomal protein L32** | GO_component: GO:0005840 - ribosome [Evidence IEA],GO_function: GO:0003735 - structural constituent of ribosome [Evidence IEA],GO_process: GO:0006412 - translation [Evidence IEA] | InterPro:IPR001515,InterPro:IPR036351,PFAM:PF01655 | note=EggNog:ENOG410PNPG,COG:J |

| | | | | | | | |
|---|---|---|---|---|---|---|---|
| chr5 | 2784402 | 2785147 | - | **ATP synthase d subunit** | GO_component: GO:0000276 - mitochondrial proton-transporting ATP synthase complex, coupling factor F(o) [Evidence IEA],GO_function: GO:0015078 - proton transmembrane transporter activity [Evidence IEA],GO_process: GO:0015986 - ATP synthesis coupled proton transport [Evidence IEA] | InterPro:IPR008689,InterPro:IPR036228,PFAM:PF05873 | note=EggNog:ENOG410PN6K,COG:C |
| chr7 | 781789 | 784130 | - | **bifunctional tryptophan synthase trp1** | GO_function: GO:0003824 - catalytic activity [Evidence IEA],GO_function: GO:0004640 - phosphoribosylanthranilate isomerase activity [Evidence IEA],GO_function: GO:0004425 - indole-3-glycerol-phosphate synthase activity [Evidence IEA],GO_function: GO:0004049 - anthranilate synthase activity [Evidence IEA],GO_process: GO:0006568 - tryptophan metabolic process [Evidence IEA] | InterPro:IPR001240,InterPro:IPR001468,InterPro:IPR006221,InterPro:IPR011060,InterPro:IPR013785,InterPro:IPR013798,InterPro:IPR016302,InterPro:IPR017926,InterPro:IPR029062, PFAM:PF00117,PFAM:PF00218,PFAM:PF00697 | note=BUSCO:EOG09261OLD,MEROPS:MER0045094,EggNog:ENOG410PGPM,COG:E |
| chr3 | 2368442 | 2369399 | + | **diphthine synthase** | GO_function: GO:0004164 - diphthine synthase activity [Evidence IEA],GO_function: GO:0008168 - methyltransferase activity [Evidence IEA],GO_process: GO:0017183 - peptidyl-diphthamide biosynthetic process from peptidyl-histidine [Evidence IEA] | InterPro:IPR000878,InterPro:IPR004551,InterPro:IPR014776,InterPro:IPR014777,InterPro:IPR035996,PFAM:PF00590 | note=BUSCO:EOG09263JW5,EggNog:ENOG410PGKF,COG:J |
| chr3 | 4044603 | 4045909 | - | **Elongation of fatty acids protein 2** | GO_function: GO:0004518 - nuclease activity [Evidence IEA],GO_function: GO:0003677 - DNA binding [Evidence IEA],GO_function: GO:0003824 - catalytic activity [Evidence IEA],GO_function: GO:0016788 - hydrolase activity, acting on ester bonds [Evidence IEA],GO_process: GO:0006281 - DNA repair [Evidence IEA] | InterPro:IPR006084,InterPro:IPR006085,InterPro:IPR006086,InterPro:IPR008918,InterPro:IPR019974,InterPro:IPR023426,InterPro:IPR029060, InterPro:IPR036279,PFAM:PF00752,PFAM:PF00867 | note=BUSCO:EOG092634B1,EggNog:ENOG410PFFR,COG:L |
| chr8 | 662079 | 663125 | + | **Eukaryotic translation initiation factor 6** | GO_function: GO:0043022 - ribosome binding [Evidence IEA],GO_process: GO:0042256 - mature ribosome assembly [Evidence IEA] | InterPro:IPR002769,PFAM:PF01912 | note=BUSCO:EOG092644O2,EggNog:ENOG410PFYZ,COG:J |
| chr1 | 2503148 | 2504115 | + | **GTP-binding nuclear protein gsp1/Ran** | GO_function: GO:0005525 - GTP binding [Evidence IEA],GO_function: GO:0003924 - GTPase activity [Evidence IEA],GO_process: GO:0006913 - nucleocytoplasmic transport [Evidence IEA] | InterPro:IPR001806,InterPro:IPR002041,InterPro:IPR005225,InterPro:IPR027417,PFAM:PF00025,PFAM:PF00071,PFAM:PF08477 | note=BUSCO:EOG09264LKR,EggNog:ENOG410PFRI,COG:U |

| | | | | | | | |
|---|---|---|---|---|---|---|---|
| chr7 | 770316 | 771581 | + | **guanine nucleotide-binding protein subunit alpha** | GO_component: GO:0005834 - heterotrimeric G-protein complex [Evidence IEA],GO_function: GO:0003924 - GTPase activity [Evidence IEA],GO_function: GO:0031683 - G-protein beta/gamma-subunit complex binding [Evidence IEA],GO_function: GO:0019001 - guanyl nucleotide binding [Evidence IEA],GO_function: GO:0005525 - GTP binding [Evidence IEA],GO_function: GO:0001664 - G protein-coupled receptor binding [Evidence IEA],GO_process: GO:0007186 - G protein-coupled receptor signaling pathway [Evidence IEA],GO_process: GO:0007165 - signal transduction [Evidence IEA] | InterPro:IPR001019,InterPro:IPR002975,InterPro:IPR011025,InterPro:IPR027417,PFAM:PF00025,PFAM:PF00503 | note=EggNog:ENOG410PHS2,COG:D,T |
| chr2 | 3857226 | 3857813 | + | **histone H2A** | GO_component: GO:0005634 - nucleus [Evidence IEA],GO_component: GO:0000786 - nucleosome [Evidence IEA],GO_function: GO:0003677 - DNA binding [Evidence IEA],GO_function: GO:0046982 - protein heterodimerization activity [Evidence IEA] | InterPro:IPR002119,InterPro:IPR007125,InterPro:IPR009072,InterPro:IPR032454,InterPro:IPR032458,PFAM:PF00125,PFAM:PF00808,PFAM:PF16211 | note=EggNog:ENOG410PNTK,COG:B |
| chr1 | 1603192 | 1603548 | + | **hypothetical protein** | | | note=EggNog:ENOG410QA9F (Taxomomic profile: 100% Hypocreales, 7 species) |
| chr1 | 2753059 | 2753857 | + | **hypothetical protein** | GO_component: GO:0000139 - Golgi membrane [Evidence IEA],GO_component: GO:0016021 - integral component of membrane [Evidence IEA],GO_component: GO:0005801 - cis-Golgi network [Evidence IEA],GO_process: GO:0006888 - endoplasmic reticulum to Golgi vesicle-mediated transport [Evidence IEA] | InterPro:IPR023601,PFAM:PF12352 | note=BUSCO:EOG09265AL8,EggNog:ENOG410PM03,COG:U |
| chr1 | 4079298 | 4080111 | - | **hypothetical protein** | GO_component: GO:0089701 - U2AF [Evidence IEA],GO_function: GO:0003723 - RNA binding [Evidence IEA],GO_function: GO:0046872 - metal ion binding [Evidence IEA],GO_function: GO:0003676 - nucleic acid binding [Evidence IEA],GO_process: GO:0000398 - mRNA splicing, via spliceosome [Evidence IEA] | InterPro:IPR000504,InterPro:IPR000571,InterPro:IPR009145,InterPro:IPR012677,InterPro:IPR035979,PFAM:PF00642 | note=EggNog:ENOG410PHGA,COG:A |

| | | | | | | | |
|---|---|---|---|---|---|---|---|
| chr1 | 4414522 | 4415168 | + | **hypothetical protein** | GO_component: GO:0005840 - ribosome [Evidence IEA],GO_function: GO:0003735 - structural constituent of ribosome [Evidence IEA],GO_function: GO:0003723 - RNA binding [Evidence IEA],GO_function: GO:0003676 - nucleic acid binding [Evidence IEA],GO_process: GO:0006412 - translation [Evidence IEA] | InterPro:IPR001892,InterPro:IPR010979,InterPro:IPR018269,InterPro:IPR027437,PFAM:PF00416 | note=BUSCO:EOG09264YEG,EggNog:ENOG410PG18,COG:J |
| chr1 | 5003097 | 5004242 | + | **hypothetical protein** | GO_function: GO:0016787 - hydrolase activity [Evidence IEA] | InterPro:IPR006680,InterPro:IPR032466,PFAM:PF04909 | note=EggNog:ENOG410PH41,COG:S |
| chr1 | 5042192 | 5042930 | - | **hypothetical protein** | GO_component: GO:0016021 - integral component of membrane [Evidence IEA] | InterPro:IPR004932,PFAM:PF03248 | note=BUSCO:EOG09265ANI,EggNog:ENOG410PM5F,COG:U |
| chr10 | 1412654 | 1413641 | - | **hypothetical protein** | GO_component: GO:0005737 - cytoplasm [Evidence IEA],GO_function: GO:0005515 - protein binding [Evidence IEA],GO_process: GO:0007015 - actin filament organization [Evidence IEA] | InterPro:IPR003005,InterPro:IPR004148,InterPro:IPR027267,InterPro:IPR037429,PFAM:PF03114 | note=EggNog:ENOG410QE92,COG:U |
| chr2 | 482295 | 483598 | - | **hypothetical protein (Actin-related protein)** | | InterPro:IPR004000,InterPro:IPR020902,PFAM:PF00022 | note=EggNog:ENOG410PHDF,COG:Z |
| chr2 | 2132703 | 2133386 | + | **hypothetical protein** | GO_component: GO:0005840 - ribosome [Evidence IEA],GO_function: GO:0003735 - structural constituent of ribosome [Evidence IEA],GO_process: GO:0006412 - translation [Evidence IEA] | InterPro:IPR001047,InterPro:IPR022309,PFAM:PF01201 | note=EggNog:ENOG410PI2H,COG:J |
| chr2 | 3027726 | 3028871 | + | **hypothetical protein** | | InterPro:IPR022057,PFAM:PF12271 | note=EggNog:ENOG410PIP5,COG:S |
| chr2 | 3283071 | 3285575 | - | **hypothetical protein** | GO_component: GO:0016020 - membrane [Evidence IEA],GO_function: GO:0016491 - oxidoreductase activity [Evidence IEA],GO_function: GO:0016651 - oxidoreductase activity, acting on NAD(P)H [Evidence IEA],GO_function: GO:0051536 - iron-sulfur cluster binding [Evidence IEA],GO_function: GO:0008137 - NADH dehydrogenase (ubiquinone) activity [Evidence IEA],GO_function: GO:0009055 - electron transfer activity [Evidence IEA],GO_process: GO:0055114 - oxidation-reduction process [Evidence IEA],GO_process: GO:0042773 - ATP synthesis coupled electron transport [Evidence IEA] | InterPro:IPR000283,InterPro:IPR001041,InterPro:IPR006656,InterPro:IPR006963,InterPro:IPR010228,InterPro:IPR015405,InterPro:IPR019574,InterPro:IPR036010,PFAM:PF00111,PFAM:PF09326,PFAM:PF10588,PFAM:PF13510 | note=EggNog:ENOG410PH1J,COG:C |

| | | | | | | | |
|---|---|---|---|---|---|---|---|
| chr2 | 4291835 | 4293444 | + | **hypothetical protein** | GO_component: GO:0005737 - cytoplasm [Evidence IEA],GO_component: GO:0005852 - eukaryotic translation initiation factor 3 complex [Evidence IEA],GO_function: GO:0003743 - translation initiation factor activity [Evidence IEA] | InterPro:IPR000717,InterPro:IPR019382,PFAM:PF10255 | note=EggNog:ENOG410PFZ7,COG:J |
| chr3 | 1915050 | 1916427 | - | **hypothetical protein** | GO_function: GO:0016747 - transferase activity, transferring acyl groups other than amino-acyl groups [Evidence IEA],GO_function: GO:0003824 - catalytic activity [Evidence IEA] | InterPro:IPR002155,InterPro:IPR016039,InterPro:IPR020610,InterPro:IPR020615,InterPro:IPR020616,InterPro:IPR020617,PFAM:PF00108,PFAM:PF02803 | note=EggNog:ENOG410PG5V,COG:I |
| chr3 | 3222971 | 3225937 | - | **hypothetical protein** | GO_function: GO:0008536 - Ran GTPase binding [Evidence IEA],GO_process: GO:0006886 - intracellular protein transport [Evidence IEA] | InterPro:IPR001494,InterPro:IPR011989,InterPro:IPR016024,PFAM:PF02985,PFAM:PF03810,PFAM:PF13513 | note=EggNog:ENOG410PJ5B,COG:U,Y |
| chr3 | 3906505 | 3906918 | + | **hypothetical protein** | GO_function: GO:0046982 - protein heterodimerization activity [Evidence IEA] | InterPro:IPR003958,InterPro:IPR009072,PFAM:PF00808 | note=EggNog:ENOG410PS30 |
| chr4 | 1363062 | 1363859 | + | **hypothetical protein** | | InterPro:IPR009038,InterPro:IPR036598,PFAM:PF01105 | note=EggNog:ENOG410PK0H,COG:U |
| chr4 | 1476210 | 1476771 | - | **hypothetical protein** | GO_component: GO:0030532 - small nuclear ribonucleoprotein complex [Evidence IEA],GO_process: GO:0008380 - RNA splicing [Evidence IEA] | InterPro:IPR001163,InterPro:IPR010920,InterPro:IPR027248,PFAM:PF01423 | note=BUSCO:EOG09265IT6,EggNog:ENOG410PNRK,COG:A |
| chr4 | 2371809 | 2372304 | + | **hypothetical protein** | | InterPro:IPR005651,PFAM:PF03966 | note=BUSCO:EOG09265F2Y,EggNog:ENOG410PPKG,COG:S (Taxomomic profile: 100% Dikaryote fungi, 72 species) |
| chr4 | 2539144 | 2540372 | + | **hypothetical protein** | GO_function: GO:0005515 - protein binding [Evidence IEA] | InterPro:IPR001680,InterPro:IPR015943,InterPro:IPR017986,InterPro:IPR019775,InterPro:IPR020472,InterPro:IPR036322,PFAM:PF00400 | note=EggNog:ENOG410PJ2T,COG:A |
| chr7 | 1225597 | 1228359 | + | **hypothetical protein** | GO_function: GO:0005524 - ATP binding [Evidence IEA],GO_process: GO:0019538 - protein metabolic process [Evidence IEA] | InterPro:IPR001270,InterPro:IPR003593,InterPro:IPR003959,InterPro:IPR004176,InterPro:IPR018368,InterPro:IPR019489,InterPro:IPR027417,InterPro:IPR028299,InterPro:IPR036628,InterPro:IPR041546,PFAM:PF00004,PFAM:PF00158,PFAM:PF02861,PFAM:PF07724,PFAM:PF07728,PFAM:PF10431,PFAM:PF17871 | note=EggNog:ENOG410PGGQ,COG:O |

| | | | | | | | |
|---|---|---|---|---|---|---|---|
| chr7 | 2007804 | 2008631 | - | **hypothetical protein** | GO_component: GO:0005840 - ribosome [Evidence IEA],GO_function: GO:0003735 - structural constituent of ribosome [Evidence IEA],GO_process: GO:0006412 - translation [Evidence IEA] | InterPro:IPR021131,InterPro:IPR021132,InterPro:IPR036227,PFAM:PF17135 | note=EggNog:ENOG410PMXF,COG:J |
| chr4 | 2095406 | 2097441 | - | **Importin alpha subunit (Karyopherin alpha subunit) (Serine-rich RNA polymerase I suppressor protein)** | GO_component: GO:0005634 - nucleus [Evidence IEA],GO_component: GO:0005737 - cytoplasm [Evidence IEA],GO_function: GO:0005515 - protein binding [Evidence IEA],GO_function: GO:0140142 - nucleocytoplasmic carrier activity [Evidence IEA],GO_function: GO:0061608 - nuclear import signal receptor activity [Evidence IEA],GO_process: GO:0006606 - protein import into nucleus [Evidence IEA] | InterPro:IPR000225,InterPro:IPR002652,InterPro:IPR011989,InterPro:IPR016024,InterPro:IPR024931,InterPro:IPR032413,InterPro:IPR036975,PFAM:PF00514,PFAM:PF01749,PFAM:PF02985,PFAM:PF13513,PFAM:PF13646,PFAM:PF16186 | note=BUSCO:EOG09261OSU,EggNog:ENOG410PI6Q,COG:U |
| chr1 | 1058610 | 1059442 | + | **kinetochore-associated Ndc80 complex subunit spc25** | | InterPro:IPR013255,PFAM:PF08234 | note=EggNog:ENOG410PHDK,COG:S |
| chr3 | 2787435 | 2788130 | + | **mitochondrial membrane protein** | GO_function: GO:0005515 - protein binding [Evidence IEA],GO_process: GO:0000266 - mitochondrial fission [Evidence IEA] | InterPro:IPR011990,InterPro:IPR016543,InterPro:IPR028058,InterPro:IPR028061,InterPro:IPR033745,PFAM:PF14852,PFAM:PF14853 | note=BUSCO:EOG09265A4E,EggNog:ENOG410PMVK,COG:M |
| chr1 | 3675964 | 3677978 | + | **Phosphoglucomutase-2** | GO_function: GO:0016868 - intramolecular transferase activity, phosphotransferases [Evidence IEA],GO_process: GO:0005975 - carbohydrate metabolic process [Evidence IEA],GO_process: GO:0071704 - organic substance metabolic process [Evidence IEA] | InterPro:IPR005841,InterPro:IPR005844,InterPro:IPR005845,InterPro:IPR005846,InterPro:IPR016055,InterPro:IPR036900,PFAM:PF02878,PFAM:PF02879,PFAM:PF02880 | note=EggNog:ENOG410PHM1,COG:G |
| chr3 | 2221373 | 2223768 | + | **Pre-mRNA-splicing factor cef1** | GO_function: GO:0003677 - DNA binding [Evidence IEA] | InterPro:IPR001005,InterPro:IPR009057,InterPro:IPR017930,InterPro:IPR021786,PFAM:PF00249,PFAM:PF11831,PFAM:PF13921 | note=BUSCO:EOG09261CXQ,EggNog:ENOG410PFBZ,COG:K |
| chr1 | 3739004 | 3739974 | - | **proteasome core particle subunit beta 2** | GO_component: GO:0005839 - proteasome core complex [Evidence IEA],GO_function: GO:0004175 - endopeptidase activity [Evidence IEA],GO_function: GO:0004298 - threonine-type endopeptidase activity [Evidence IEA],GO_process: GO:0051603 - proteolysis involved in cellular protein catabolic process [Evidence IEA] | InterPro:IPR000243,InterPro:IPR001353,InterPro:IPR023333,InterPro:IPR024689,InterPro:IPR029055,PFAM:PF00227,PFAM:PF12465 | note=BUSCO:EOG092641M3,MEROPS:MER0000542,EggNog:ENOG410PH45,COG:O |

| | | | | | | | |
|---|---|---|---|---|---|---|---|
| chr1 | 3802131 | 3802822 | - | **ribosomal 40S subunit protein S13** | GO_component: GO:0005840 - ribosome [Evidence IEA],GO_function: GO:0003735 - structural constituent of ribosome [Evidence IEA],GO_process: GO:0006412 - translation [Evidence IEA] | InterPro:IPR000589,InterPro:IPR009068,InterPro:IPR012606,InterPro:IPR023029,PFAM:PF00312,PFAM:PF08069 | note=BUSCO:EOG0926514P,EggNog:ENOG410PMVC,COG:J |
| chr2 | 2974410 | 2975180 | + | **ribosomal protein S5** | GO_component: GO:0015935 - small ribosomal subunit [Evidence IEA],GO_function: GO:0003735 - structural constituent of ribosome [Evidence IEA],GO_function: GO:0003723 - RNA binding [Evidence IEA],GO_process: GO:0006412 - translation [Evidence IEA] | InterPro:IPR000235,InterPro:IPR005716,InterPro:IPR020606,InterPro:IPR023798,InterPro:IPR036823,PFAM:PF00177 | note=EggNog:ENOG410PIAI,COG:J |
| chr1 | 3035216 | 3036146 | - | **RIBULOSE-phosphate 3-epimerase** | GO_function: GO:0004750 - ribulose-phosphate 3-epimerase activity [Evidence IEA],GO_function: GO:0016857 - racemase and epimerase activity, acting on carbohydrates and derivatives [Evidence IEA],GO_function: GO:0003824 - catalytic activity [Evidence IEA],GO_process: GO:0006098 - pentose-phosphate shunt [Evidence IEA],GO_process: GO:0005975 - carbohydrate metabolic process [Evidence IEA] | InterPro:IPR000056,InterPro:IPR011060,InterPro:IPR013785,InterPro:IPR026019,PFAM:PF00834 | note=BUSCO:EOG092644Z6,EggNog:ENOG410PGBP,COG:G |
| chr3 | 2136284 | 2136812 | + | **RNA polymerase II mediator complex subunit** | GO_component: GO:0016592 - mediator complex [Evidence IEA] | InterPro:IPR021384,InterPro:IPR037212,PFAM:PF11221 | note=BUSCO:EOG09265SHM,EggNog:ENOG410PQXU,COG:K |
| chr5 | 2053980 | 2054325 | - | **Sec61p translocation complex subunit** | GO_component: GO:0016020 - membrane [Evidence IEA],GO_function: GO:0015450 - P-P-bond-hydrolysis-driven protein transmembrane transporter activity [Evidence IEA],GO_process: GO:0006886 - intracellular protein transport [Evidence IEA],GO_process: GO:0006605 - protein targeting [Evidence IEA],GO_process: GO:0015031 - protein transport [Evidence IEA] | InterPro:IPR001901,InterPro:IPR008158,InterPro:IPR023391,PFAM:PF00584 | note=BUSCO:EOG09265PUI,EggNog:ENOG410PRYA,COG:U |
| chr5 | 2037555 | 2039619 | + | **Sulfate adenylyltransferase** | GO_function: GO:0004781 - sulfate adenylyltransferase (ATP) activity [Evidence IEA],GO_function: GO:0005524 - ATP binding [Evidence IEA],GO_function: GO:0004020 - adenylylsulfate kinase activity [Evidence IEA],GO_process: GO:0000103 - sulfate assimilation [Evidence IEA],GO_process: GO:0000096 - sulfur amino acid metabolic process [Evidence IEA] | InterPro:IPR002650,InterPro:IPR002891,InterPro:IPR014729,InterPro:IPR015947,InterPro:IPR024951,InterPro:IPR025980,InterPro:IPR027417,InterPro:IPR027535,PFAM:PF01583,PFAM:PF01747,PFAM:PF14306 | note=EggNog:ENOG410PFXE,COG:P |

| | | | | | | | |
|---|---|---|---|---|---|---|---|
| chr3 | 412800 | 415154 | + | **Translation initiation factor 3 subunit b** | GO_component: GO:0005852 - eukaryotic translation initiation factor 3 complex [Evidence IEA],GO_function: GO:0003743 - translation initiation factor activity [Evidence IEA],GO_function: GO:0003723 - RNA binding [Evidence IEA],GO_function: GO:0031369 - translation initiation factor binding [Evidence IEA],GO_function: GO:0005515 - protein binding [Evidence IEA],GO_function: GO:0003676 - nucleic acid binding [Evidence IEA],GO_process: GO:0006413 - translational initiation [Evidence IEA] | InterPro:IPR000504,InterPro:IPR011400,InterPro:IPR012677,InterPro:IPR013979,InterPro:IPR015943,InterPro:IPR034363,InterPro:IPR035979,PFAM:PF08662 | note=BUSCO:EOG09260K24,EggNog:ENOG410PGHS,COG:J |
| chr1 | 4128297 | 4129890 | - | **translation initiation factor eIF4A** | GO_function: GO:0005524 - ATP binding [Evidence IEA],GO_function: GO:0003676 - nucleic acid binding [Evidence IEA] | InterPro:IPR000629,InterPro:IPR001650,InterPro:IPR011545,InterPro:IPR014001,InterPro:IPR014014,InterPro:IPR027417,PFAM:PF00270,PFAM:PF00271,PFAM:PF04851 | note=EggNog:ENOG410PFHJ,COG:J |
| chr4 | 3873687 | 3877402 | + | **translational elongation factor EF-1 alpha** | GO_function: GO:0005524 - ATP binding [Evidence IEA],GO_function: GO:0016887 - ATPase activity [Evidence IEA] | InterPro:IPR003439,InterPro:IPR003593,InterPro:IPR011989,InterPro:IPR016024,InterPro:IPR017871,InterPro:IPR021133,InterPro:IPR027417,InterPro:IPR040533,PFAM:PF00005,PFAM:PF17947 | note=EggNog:ENOG410PGET,COG:J |
| chr1 | 1297736 | 1298401 | + | **Ubiquitin-conjugating enzyme E2 11** | | InterPro:IPR000608,InterPro:IPR016135,InterPro:IPR023313,PFAM:PF00179 | note=EggNog:ENOG410PNQ1,COG:O |
| chr3 | 1917440 | 1918078 | + | **Ubiquitin-conjugating enzyme E2 2** | | InterPro:IPR000608,InterPro:IPR016135,InterPro:IPR023313,PFAM:PF00179 | note=EggNog:ENOG410PHPZ,COG:K |

### Table S4 | Genome annotation summary statistics for *Neonectria neomacrospora* (KNNDK1)

| Chromosome | 1 | 2 | 3 | 4 | 5 | 6 | 7 | 8 | 9 | 10 | 11 | 12 |
|---|---|---|---|---|---|---|---|---|---|---|---|---|
| Size (Mb) | 6.42 | 5.13 | 4.82 | 4.62 | 3.57 | 3.03 | 2.50 | 2.06 | 1.98 | 1.80 | 0.87 | 0.30 |
| Size (bp) | 6,419,651 | 5,126,922 | 4,819,673 | 4,617,141 | 3,569,078 | 3,030,596 | 2,503,575 | 2,056,958 | 1,982,407 | 1,804,562 | 865,340 | 296,394 |
| GC content (%) | 54.68 | 54.46 | 54.05 | 53.32 | 53.46 | 53.14 | 53.75 | 52.97 | 51.51 | 52.77 | 48.47*** | 39.12*** |
| Number of protein-coding genes | 1,946 | 1,554 | 1,418 | 1,474 | 1,051 | 967 | 721 | 655 | 608 | 592 | 268 | 37 |
| Mean gene length (bp) | 1,649 | 1,692 | 1,760 | 1,676 | 1,687 | 1,572 | 1,790 | 1,526 | 1,604 | 1,559 | 1,581* | 1,274*** |
| Genes content (%) | 73.04 | 51.29 | 51.77 | 53.51 | 49.68 | 50.15 | 51.56 | 48.58 | 49.20 | 51.13 | 48.97 | 15.91*** |
| Proportion of genes with unknown function(%) | 5.73 | 5.21 | 5.00 | 5.02 | 6.37 | 6.51 | 4.02 | 4.12 | 6.91 | 4.73 | 8.21*** | 54.05*** |
| Signal peptide[1] | 143 | 121 | 109 | 144 | 59 | 118 | 52 | 72 | 64 | 62 | 43 | 0 |
| Of which is CAZy | 52 | 19 | 21 | 37 | 10 | 32 | 7 | 15 | 17 | 20 | 14 | 0 |
| Of which is proteases | 18 | 19 | 19 | 13 | 12 | 14 | 5 | 5 | 4 | 4 | 6 | 0 |
| Both of the above | 1 | 1 | 0 | 0 | 0 | 0 | 0 | 0 | 0 | 0 | 0 | 0 |
| CAZymes[2] | 70 | 47 | 56 | 69 | 32 | 49 | 21 | 24 | 30 | 35 | 18 | 0 |
| CAZymes per Mb | 10.90 | 8.87 | 11.62 | 14.94 | 8.96 | 16.17 | 8.40 | 11.65 | 15.15 | 19.44 | 20.69 | 0 |
| Proteases[3] | 67 | 53 | 48 | 51 | 35 | 40 | 28 | 21 | 20 | 19 | 12 | 1 |
| CAZymes-Proteases | 6 | 3 | 2 | 3 | 5 | 2 | 2 | 0 | 2 | 2 | 1 | 0 |

Asterisks indicate data for the mini-chromosomes differ significantly from the mean for chromosomes 1-10 (One-Sample t-test. *** P <0.001; ** P <0.01; * P <0.05). For signal peptides. CAZymes and proteases are all numbers given counts.

[1] Signal peptides predicted with SignalP-5.

[2] Annotated using dbCAN

[3] Annotated with MEROPS
